# Interaction Profiles as a Universal Language for Generative Molecular Design with ShEPhERD–2

**DOI:** 10.64898/2026.09.10.750648

**Authors:** Kento A. Abeywardane, Kenji Walker, Connor W. Coley

## Abstract

Three-dimensional intermolecular interactions govern molecular recognition and are fundamental to the pharmacological activity of small-molecule drugs. We propose that an interaction profile, comprising shape, electrostatics, and directional pharmacophores, is a sufficient and transferable design specification for molecular design. We introduce ShEPhERD–2, a 3D generative model that generates molecular structures conditioned on explicit interaction profiles. ShEPhERD–2 generates low-strain, drug-like molecules more efficiently and with greater fidelity to target interaction profiles than its predecessor while enabling precise control over molecular design through pharmacophore prioritization, substructure constraints, and composition of multiple interaction profiles. We demonstrate the application of a single model to bioisosteric fragment merging, dual-target design, selectivity engineering, and modality hopping without task-specific retraining. These results establish interaction profiles as a chemotype-agnostic interface between structure hypotheses and molecular design.

## 1 Introduction

The three-dimensional interactions between a ligand and its targets govern molecular recognition, dictating potency, selectivity, and other key pharmacological properties [1, 2]. Accordingly, a principal aim of medicinal chemistry is to identify the interactions driving on- and off-target recognition and then to design molecules that satisfy the desired interaction pattern [3]. While this aim is fundamental to drug design, it requires navigating a vast chemical and conformational space subject to complex chemical and geometric constraints. This difficulty has motivated molecular generative models that, in principle, directly generate 3D structures meeting these constraints.

Most current 3D generative models for small-molecule design generate all-atom molecular structures under the implicit assumption that favorable interactions, and the appropriate chemical structure that satisfies them, can be learned end-to-end from geometric patterns in their training data [4–7]. For example, a model may learn from protein–ligand complexes that an amide N-H and a carbonyl O are frequently found ~ 2.9 Å apart, without relating this to the generalizable notion of a hydrogen bond [8]. While appealing in theory, these approaches have two limitations for small molecule design: (1) they must infer all principles of molecular recognition from a limited corpus of experimentally determined protein–ligand complexes or from approximate *simulated* poses, and, as a result, often fail to recapitulate native interactions, adopt strained conformations, and over-rely on nonselective hydrophobic contacts [9–12]; and (2) because interactions are never explicit design variables, medicinal chemists cannot directly incorporate prior knowledge of desired interactions.

Aligning generative models more closely with the intuition of medicinal chemists through explicit interaction representations, grounded in decades of cheminformatics research [13–19], offers a promising route to overcoming these limitations. This has motivated the nascent paradigm of *interaction-aware* generative models, which enable medicinal chemists to specify desired interaction hypotheses directly. Such hypotheses can be extracted from varied experimental discovery sources, such as ligand conformers, bound fragments, or native biomolecules, and can be realized by diverse molecules, providing a versatile input representation for generative modeling across small molecule discovery [20, 21]. Moreover, because potential interactions can be derived from ligand conformers alone, there is an opportunity to leverage abundant ligand-only 3D structural data. Existing interaction-aware models have explored diverse facets of molecular recognition with different levels of detail, including molecular shape [22–26], pharmacophores [27–29] and specific non-covalent interactions [30–34]. However, these models typically focus on subsets of interactions, often omit considerations of directionality, and provide limited control over how interactions are satisfied, leaving opportunities to improve the versatility with which interaction design objectives can be expressed and refined. Concurrent with and complementary to our work, Irwin et al. [35] explore improving the versatility of interaction-aware models through additional conditioning channels to encode conformer ensemble properties; our work focuses on unifying and controlling how interactions condition molecular generation.

In this study, we introduce ShEPhERD–2^1^, a 3D generative model that enables flexible specification, prioritization, and composition of molecular interaction patterns for interaction-driven design of small molecules. This framework is centered on the *interaction profile*: a joint representation of solvent-accessible surface, electrostatic potential, and pharmacophoric features (Fig. 1a) that captures key geometric and physicochemical determinants of molecular recognition, introduced in our prior work ShEPhERD [36]. ShEPhERD–2 uses an SE(3)-equivariant diffusion framework to jointly generate 3D molecular structures and their complete interaction profiles (Fig. 1b), achieving higher conformer quality, improved generation efficiency, and more faithful recapitulation of target interactions when conditioned on a query interaction profile than its predecessor. Building on this foundation, we introduce generative capabilities to prioritize subsets of pharmacophoric interactions, constrain molecular substructures or scaffolds, and compose multiple on- and off-target interaction profiles. Together, these capabilities enable a broad range of small-molecule design applications, including hit expansion, scaffold elaboration, interaction-space optimization, dual-target design, selectivity engineering, and modality-hopping tasks such as peptide mimicry (Fig. 1c-h). Demonstrations of ShEPhERD–2’s application to these diverse tasks illustrate how interaction profiles can serve as a universal language for generative molecular design.

**Fig. 1:**
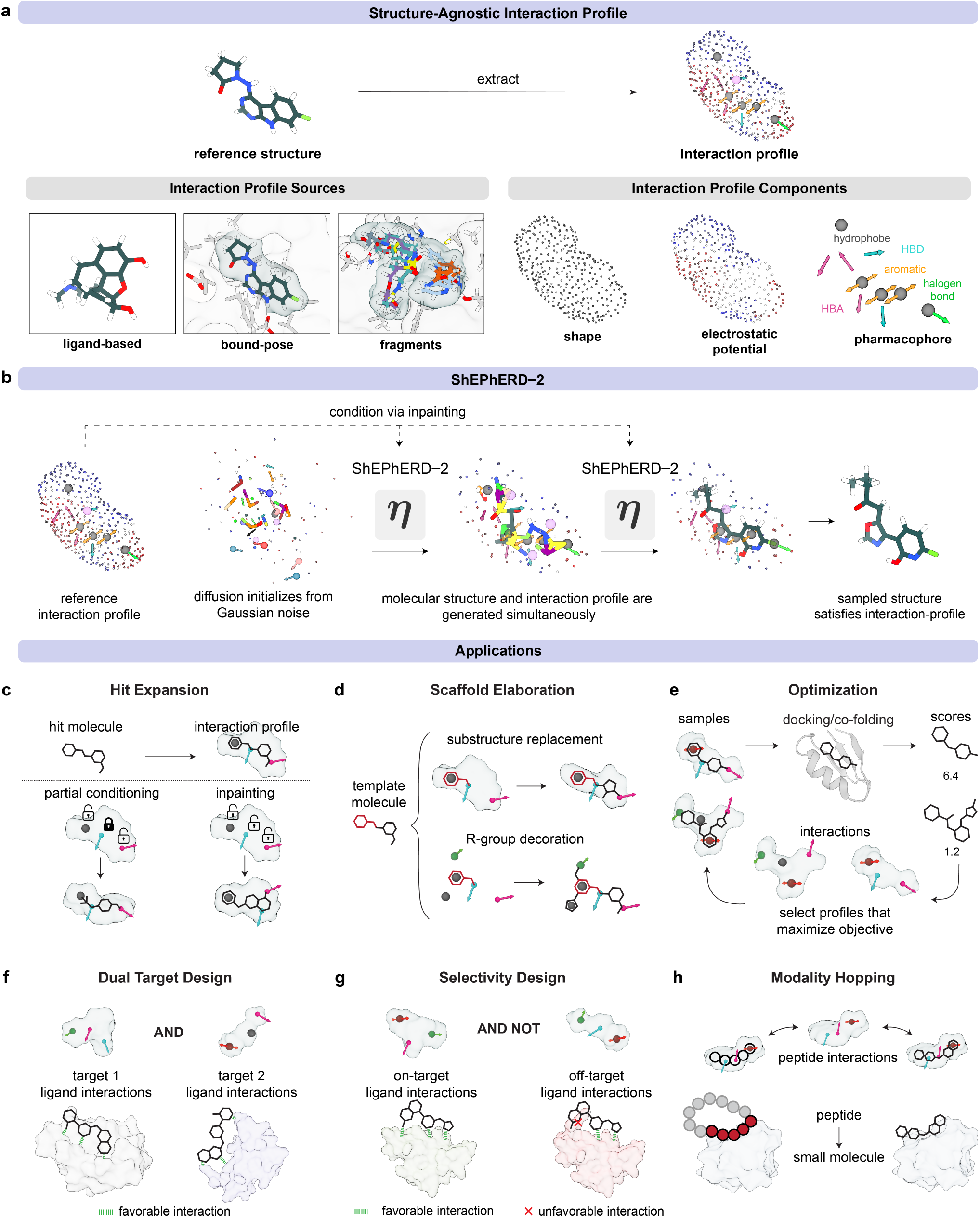
**a**, Chemical structures sourced from various discovery modalities can be abstracted into a common interaction profile comprising molecular shape, electrostatic potential, and pharmacophoric features. **b**, ShEPhERD–2 simultaneously denoises molecular structure and interaction profiles. Interaction profiles extracted from reference molecules are used as conditioning signals via inpainting, enabling generation of chemically diverse molecules that preserve desired interaction patterns via denoising diffusion. **c**, Interaction-conditioned hit expansion generates alternative chemotypes while maintaining key interactions in a flexible manner. **d**, Fixed-atom constraints enable scaffold elaboration or growth around user-specified substructures. **e**, Interaction profiles can be optimized directly with an evolutionary algorithm to improve molecular recognition as evaluated by a computational oracle. **f**, Multiple interaction profiles can be composed to generate ligands that satisfy the interaction requirements of two targets simultaneously. **g**, Interaction profile composition can also promote target selectivity by optimizing one interaction profile while avoiding another. **h**, Because interaction profiles provide a modality-independent representation of molecular recognition, ShEPhERD–2 enables modality hopping, such as designing small molecules that mimic peptide interactions.

## 2 Results

### 2.1 Interaction-guided generation for analoging and hit expansion

A central challenge in interaction-driven design is to identify novel molecules with realistic conformations that exhibit a desired three-dimensional interaction profile. ShEPhERD was previously introduced as an interaction-aware generative diffusion framework that jointly generates molecular structure, solvent-accessible surface, electrostatic potential (ESP), and pharmacophoric features [36]. ShEPhERD–2 extends this framework with methodological advances that improve sampling robustness, conformational realism, and computational efficiency. Trained on 1.6 million GFN2-xTB [37] locally relaxed drug-like molecules from the MOSES dataset [36, 38], ShEPhERD– 2 learns exclusively from interaction profiles computed directly from molecular geometry, eliminating the need for protein–ligand complex data and enabling training on large-scale virtually enumerated libraries.

ShEPhERD–2 allows for the generation of molecules of different sizes without *a priori* specification via dummy atoms that let superfluous atoms be removed during denoising [39, 40]. This expands the set of molecules accessible from a given initialization, enabling the model to converge to an appropriate molecular size while preserving chemically plausible hybridization (Fig. 2a; Suppl. Fig. S1a). ShEPhERD– 2 further replaces the original diffusion process with a variance-exploding formulation and adopts a more efficient SE(3)-equivariant architecture that reduces conformational strain of its proposed geometries and, with other optimizations, accelerates sampling by 6.4× (Suppl. Table S5-S6; Suppl. Fig. S1b).

**Fig. 2:**
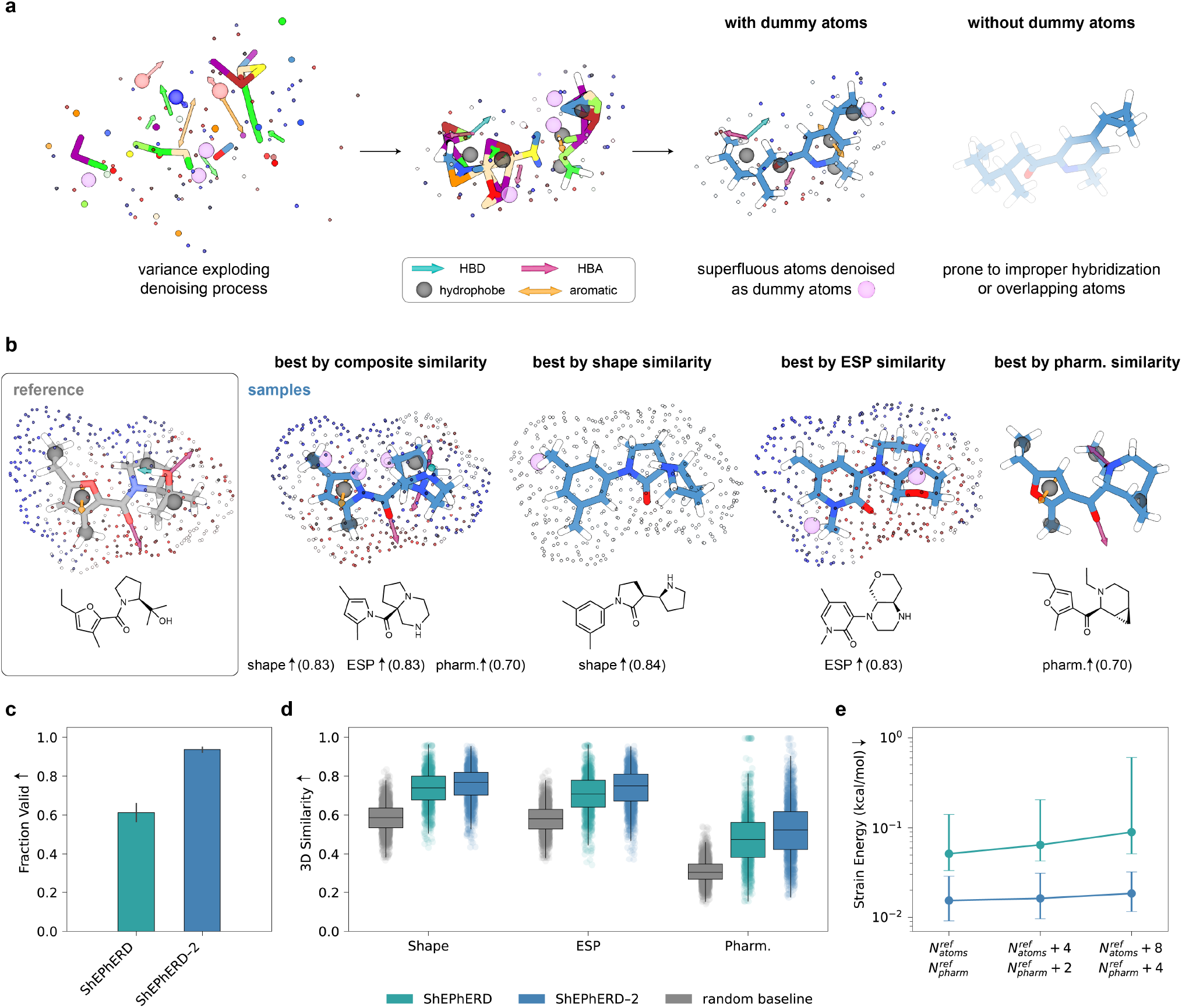
**a**, Representative variance-exploding denoising trajectory illustrating the role of absorbing dummy atoms (pink). By allowing superfluous atoms to transition into dummy states, ShEPhERD–2 expands the set of molecules accessible from a given initialization. Without dummy atoms, the same initialization yields a molecule with improper hybridization. **b**, Representative molecules generated from conditioning on the full interaction profile of a reference molecule. From graph-diverse samples (fingerprint similarity to reference *<*0.3), we selected molecules with high composite interaction similarity (shape + ESP + pharmacophore) or high similarity in each individual interaction modality. 3D similarity is computed after GFN2-xTB relaxation and re-alignment. We show only the sample interaction profile components corresponding to the reported similarity scores. Surface representations are upsampled for visual clarity. **c**, ShEPhERD–2 achieves high validity under interaction-conditioned generation. **d**, ShEPhERD–2 recapitulates target shape, electrostatic potential (ESP), and pharmacophoric features more faithfully than both a random baseline and ShEPhERD. **e**, ShEPhERD–2 maintains low conformational strain across increasing atom-count initializations, whereas ShEPhERD produces progressively more strained conformations.

Interaction-conditioned generation is achieved using inpainting [41–44], which stochastically guides the diffusion trajectory to reconstruct a specified interaction profile. This allows the model to generate the molecular structure freely, yielding chemically distinct molecules that recapitulate the interaction profile. Representative samples have high 3D interaction similarity for the complete interaction profile and for individual interaction modalities, while achieving low graph similarity (*<* 0.3) to the reference (Fig. 2b).

Quantitative evaluations show significant improvement in sample quality over the original ShEPhERD model. Across 100 molecules from the MOSES scaffold-split test set [38], we generated 20 candidates conditioned on each target interaction profile. ShEPhERD–2 produced chemically valid molecules 93.7% of the time, compared with 61.3% for the original ShEPhERD framework (Fig. 2c; Suppl. Table S7). Among graph-diverse samples, ShEPhERD–2 more faithfully reproduced the target interaction profiles, yielding higher average and top-1 shape, electrostatic potential, and pharmacophore similarity (all Mann–Whitney *p <* 10^−4^; Fig. 2d; Suppl. Fig. S2). These improvements were accompanied by low conformational strain following local GFN2-xTB relaxation, which remained nearly constant even when generation was initialized with excess atoms and pharmacophoric features (Fig. 2e), demonstrating robust performance across a broad range of initialization states. Together, these advances enable ShEPhERD–2 to generate chemically valid, low-strain molecules that more faithfully reproduce target interaction profiles, establishing a robust foundation for the controllable interaction-guided design tasks explored below.

### 2.2 Controllable molecular generation with pharmacophore and scaffold constraints

Not all protein–ligand interactions are created equal. Certain pharmacophores may require precise positioning and orientation to maintain critical contacts with protein residues, whereas others, such as solvent-exposed groups or nonspecific hydrophobic interactions, can be relaxed. Different types of interactions might also contribute more or less to the overall energetics of binding [45–47]. Likewise, preserving a known scaffold may be desirable because it confers favorable physicochemical properties or synthetic accessibility. ShEPhERD–2 enables both capabilities by fixing selected pharmacophores or substructures.

For pharmacophore prioritization, selected high-priority (HP) pharmacophores remain fixed throughout the generative diffusion trajectory, whereas low-priority (LP) pharmacophores are generated using standard, stochastic inpainting. This enables users to tune the fidelity with which different interactions are realized, enforcing critical pharmacophores while maintaining flexibility elsewhere in the molecule. Fig. 3a illustrates this behavior with a representative example. The aromatic pharmacophore of the fluorobenzene ring and the hydrogen-bond acceptor of the 2-fluoropyridine were designated as HP and remained geometrically conserved after GFN2-xTB relaxation of the generated samples, while the remaining structures exhibited greater diversity in LP pharmacophoric feature placement and produced more structurally diverse motifs.

**Fig. 3:**
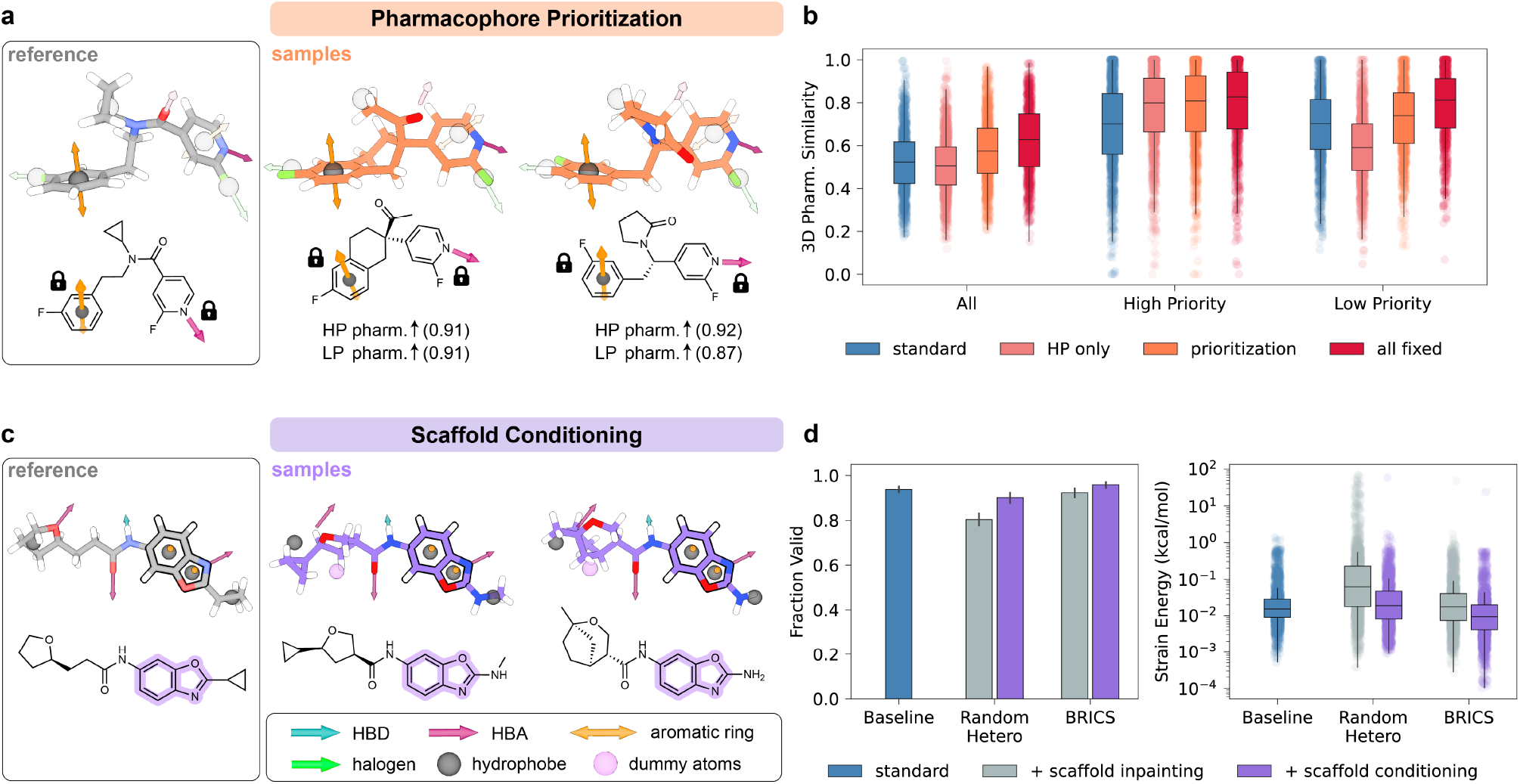
**a**, High-priority (HP) pharmacophores are highlighted on the reference structure (gray), and the remaining, transparent features are designated as low priority (LP). Generated molecules (orange) preserve the HP pharmacophore geometry with high fidelity while LP features are retained but allowed greater geometric flexibility as shown when overlaid with the reference molecule’s pharmacophore. **b**, Distributions of 3D pharmacophore similarity for the four generation strategies. *Standard* inpaints all pharmacophores. *HP only* fixes the HP pharmacophore while leaving LP pharmacophores unconstrained. *Prioritization* fixes the HP pharmacophore and inpaints LP pharmacophores. *All fixed* fixes both HP and LP pharmacophores throughout generation. Similarity is computed relative to the reference subset specified by each strategy. **c**, The BRICS-identified benzoxazole scaffold of the reference molecule (gray) that is fixed during generation is highlighted on the reference chemical structure. ShEPhERD–2 samples (purple), overlaid with the reference molecule’s pharmacophore, contain this scaffold with diversified substituents and preserved interactions. **d**, Validity and strain energy when fixing random heteroatoms or a BRICS scaffold are maintained relative to standard interaction-conditioned sampling and out-perform scaffold inpainting.

We evaluated prioritization across the full MOSES test set by randomly assigning a subset of pharmacophores as high-priority and comparing four conditioning strategies: standard inpainting, fixing only the high-priority pharmacophore, prioritization (fixed HP with LP inpainting), and fixing all pharmacophoric features. Conditioning behavior followed the expected trend: constraining additional pharmacophoric features progressively increased recovery of the complete pharmacophore profile, with fixing all features achieving the highest similarity (median 0.627). Evaluating pharmacophore similarity after relaxation prevents perfect recovery when all pharmacophores are fixed. Prioritization selectively improved recovery of high-priority pharmacophores (median 0.808 versus 0.701 for standard inpainting; *p <* 10^−4^) while maintaining flexibility in the unconstrained features. In contrast, fixing only the high-priority subset reduced recovery of low-priority pharmacophores (median 0.590), whereas prioritization preserved substantially higher recovery (median 0.739; *p <* 10^−4^). These results demonstrate that ShEPhERD–2 enables flexible control over which interaction features are enforced during generation (Fig. 3a; Suppl. Table S9).

We next applied the same conditioning strategy to scaffold-constrained generation by fixing selected atomic coordinates and identities throughout denoising rather than reconstructing them through stochastic inpainting [6, 48]. This avoids the strained conformations introduced by inpainting and reduces steric clashes during generation. Representative examples demonstrate that fixed scaffolds can be elaborated with diverse substituents while preserving the desired interaction profile. For instance, Fig. 3c shows that the fixed benzoxazole scaffold is accommodated without introducing strain into the surrounding substituents, resulting in high 3D pharmacophore similarity even after GFN2-xTB relaxation. When fixing BRICS fragments [49] or specific heteroatoms, scaffold conditioning maintained strain energies and chemical validity comparable to unconstrained generation, whereas scaffold inpainting substantially increased strain and, in the fixed-heteroatom setting, reduced validity (Fig. 3d; Suppl. Table S10-S11).

### 2.3 Interaction-guided hit-to-lead optimization without pocket structures

The pursuit of diverse chemical series during the early stages of molecular optimization can help derisk lead development by providing alternative chemotypes that preserve key intermolecular interactions but differ in properties such as synthetic accessibility, developability, and intellectual property [50]. We therefore evaluate ShEPhERD–2 on MolGenBench [51], which comprises 120 protein targets, each represented by five congeneric series of experimentally validated ChEMBL actives. In addition to the benchmark’s molecular quality and interaction metrics, we consider whether generated molecules maintain diversity across alternative chemotypes. We quantify this using two complementary measures: *Reference Tanimoto*, which measures fingerprint similarity to a reference ligand, and *Interaction Diversity*, which measures the diversity of substructures mediating conserved protein interactions across each series.

We evaluate ShEPhERD–2 on the hit-to-lead task of MolGenBench for both ligand-based and structure-based settings, and compare against representative 3D diffusion-based, interaction- and pocket-conditioned generative models, including FLOWR.ROOT [24], PhoreGen [27], OMTRA [28], CondSemla [52, 53], DiffDec [54], and DiffSBDD [6]. In the ligand-based setting, only the bound pose of the reference ligand is used to condition each generative model; this is an optimistic, best-case scenario for ligand-based design, as bound poses may be unknown in practice. The structure-based setting additionally makes use of the 3D atomic positions of pocket residues to derive interaction features. One representative, chemically distinct ligand from each series serves as the reference, and 200 candidate molecules are generated for each of the 600 references. Generated samples are assessed using MolGenBench’s suite of molecular validity, conformational quality, and structure-based interaction metrics.

ShEPhERD–2 ranks first or second across nearly every evaluation metric among ligand-based methods, providing the strongest overall balance between molecular quality, interaction preservation, docking performance, and scaffold exploration (Table 1). Reference methods FLOWR.ROOT, PhoreGen, and OMTRA are conditioned on pharmacophoric representations derived from the ligand bound pose, as is ShEPhERD–2, while CondSemla conditions directly on the reference ligand conformer. Although OMTRA matches ShEPhERD–2 in interaction recovery and docking metrics, its generated molecules remain substantially closer to the reference ligand (0.451 versus 0.199 mean Tanimoto) and exhibit markedly lower interaction diversity (0.417 versus 0.756), indicating that these interactions are often recovered by preserving the original chemotype. In contrast, ShEPhERD–2 identifies chemically distinct molecules that satisfy similar interaction constraints through alternative substructures, yielding strong interaction recovery without sacrificing scaffold novelty.

**Table 1:**
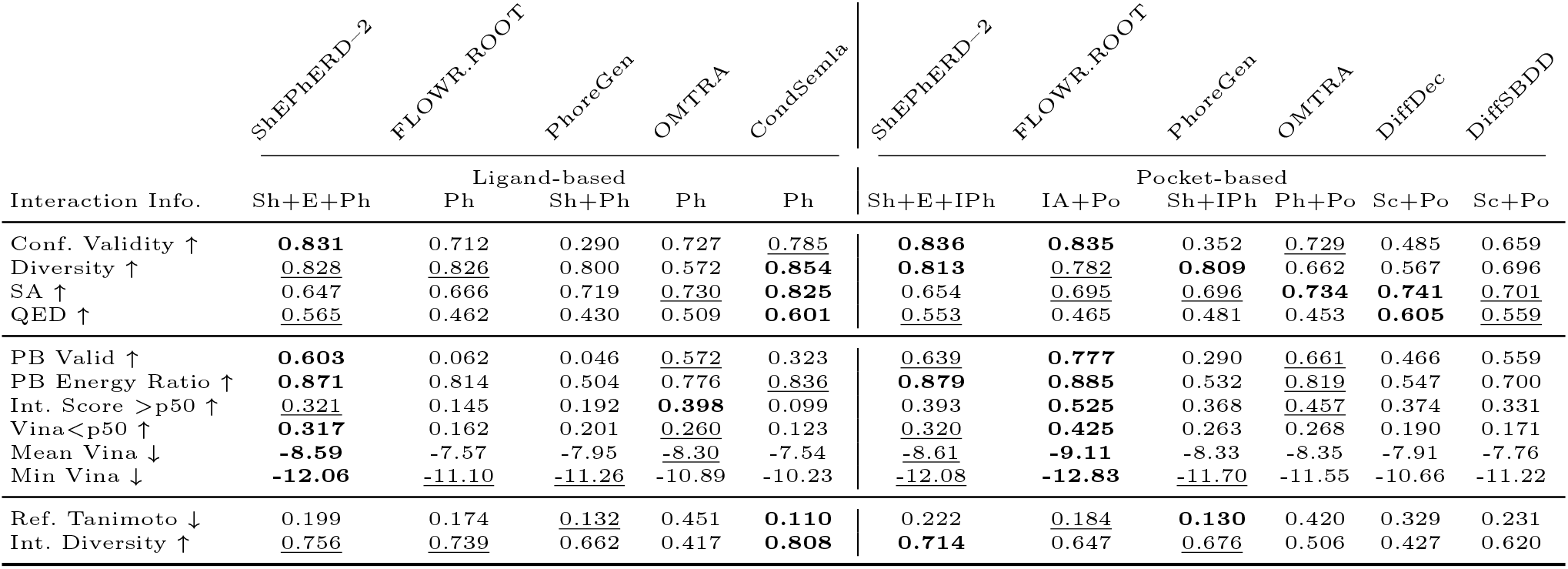
Performance on the MolGenBench hit-to-lead benchmark. Ligand-based models are conditioned only on the reference ligand, whereas pocket-based models additionally use protein pocket information when available. Conditioning modalities are summarized as combinations of molecular shape (Sh), electrostatics (E), pharmacophoric features (Ph), pocket atoms (Po), interacting pharmacophores (IPh), interacting atoms (IA), and conserved scaffold (Sc). Metrics assess molecular quality, interaction recovery, docking performance, and diversity (see Sec. 4.5.2 and Suppl. Table S17 for definitions). Best and second-best performances are shown in **bold** and <u>underlined</u>, respectively. Models that are statistically indistinguishable share a rank. Ranks come from two-sided paired tests on 10,000 cluster-bootstrap resamples of the protein targets, with family-wise error controlled at *α* = 0.05 across all pairwise comparisons within a row by step-down max-*t* (Suppl. Tables S15–S16).

We next investigated whether interaction-centric representations could recover the performance of models that explicitly encode the protein pocket. Among the structure-based baselines, FLOWR.ROOT directly conditions on both pocket geometry and ProLIF-derived interacting atoms, while OMTRA combines explicit pocket geometry with ligand-derived pharmacophores, and PhoreGen conditions on pocket-derived pharmacophores using AncPhore [55]. ShEPhERD–2 uses ligand-derived interaction profiles but prioritizes pharmacophoric features that participate in ProLIF-identified interactions. FLOWR.ROOT achieves the strongest overall performance in the pocket-based setting, leading in pocket complementarity, interaction recovery, and docking metrics (Table 1). These results are confounded by the presence of 98/120 and 70/120 of the MolGenBench PDB structures in the PLINDER [56] and CrossDocked2020 [57] training datasets used by each of these pocket-based models, so we cannot rule out the possibility that this performance is artificially inflated. Nevertheless, ShEPhERD–2 remains competitive with FLOWR.ROOT on docking performance despite not conditioning on pocket geometry directly, while holistically outperforming PhoreGen and OMTRA which struggle with conformer validity and diversity, respectively.

ShEPhERD–2 leads all models in interaction diversity with sufficient dissimilarity to the reference molecule. These results indicate that abstract, prioritized pharmacophoric constraints recover much of the information required for pocket-aware hit diversification while encouraging exploration of alternative interaction motifs rather than reproducing the reference chemotype. Consistent with this interpretation, replacing pharmacophore prioritization with direct conditioning on interacting atoms recovers most of FLOWR.ROOT’s affinity advantage (Interaction Score *>*p50: 0.514 vs. 0.525; mean Vina: −8.66 vs. −9.11 kcal/mol), albeit at the expense of PoseBusters validity and diversity (Suppl. Table S16). This indicates that the reference interactions account for most of the achievable gains, while explicit pocket geometry provides comparatively modest improvements, corroborating the central premise of this work.

### 2.4 Flexible molecular generation via composition of on- and off-target interaction profiles

While the consideration of molecular interactions for on-target potency may be of primary importance, selectivity and polypharmacology can also be engineered by considering interactions across multiple targets simultaneously. ShEPhERD–2 supports this multi-target design at inference-time through compositional logic across interaction profiles. Two forms of pairwise composition are considered: AND composition, which steers generation toward molecules that satisfy two interaction profiles simultaneously, and NOT composition, which steers generation toward molecules that satisfy one interaction profile while avoiding another. We extend a compositional diffusion approach developed for image synthesis [58], where each condition’s model output is linearly combined at every denoising step to sample from a product or negation distribution (Sec. 4.2).

Representative examples in Fig. 4a illustrate both AND and NOT composition behavior. In the AND composition sample, the amide hydrogen bond donor of **1** is combined with the furan of **2** to yield a pyrrole. In doing so, the generated molecule finds a compromise solution between the hydrogen bond donor from profile **1** and the aromatic group from profile **2**, with only a small deviation from the desired positioning of the aromatic ring pharmacophore. For NOT composition, the generated molecule achieves dissimilarity through two mechanisms. First, revision of the central ester in **1** to a carbonate ester shifts the position of a tetrahydroindolizine ring, allowing a methyl hydrophobe to satisfy **1**’s cyclopentane hydrophobe while remaining farther from **2**’s trifluoromethyl hydrophobe. Second, the fused ring reduces shape complementarity to the molecule **2** by extending outside its solvent-accessible surface.

**Fig. 4:**
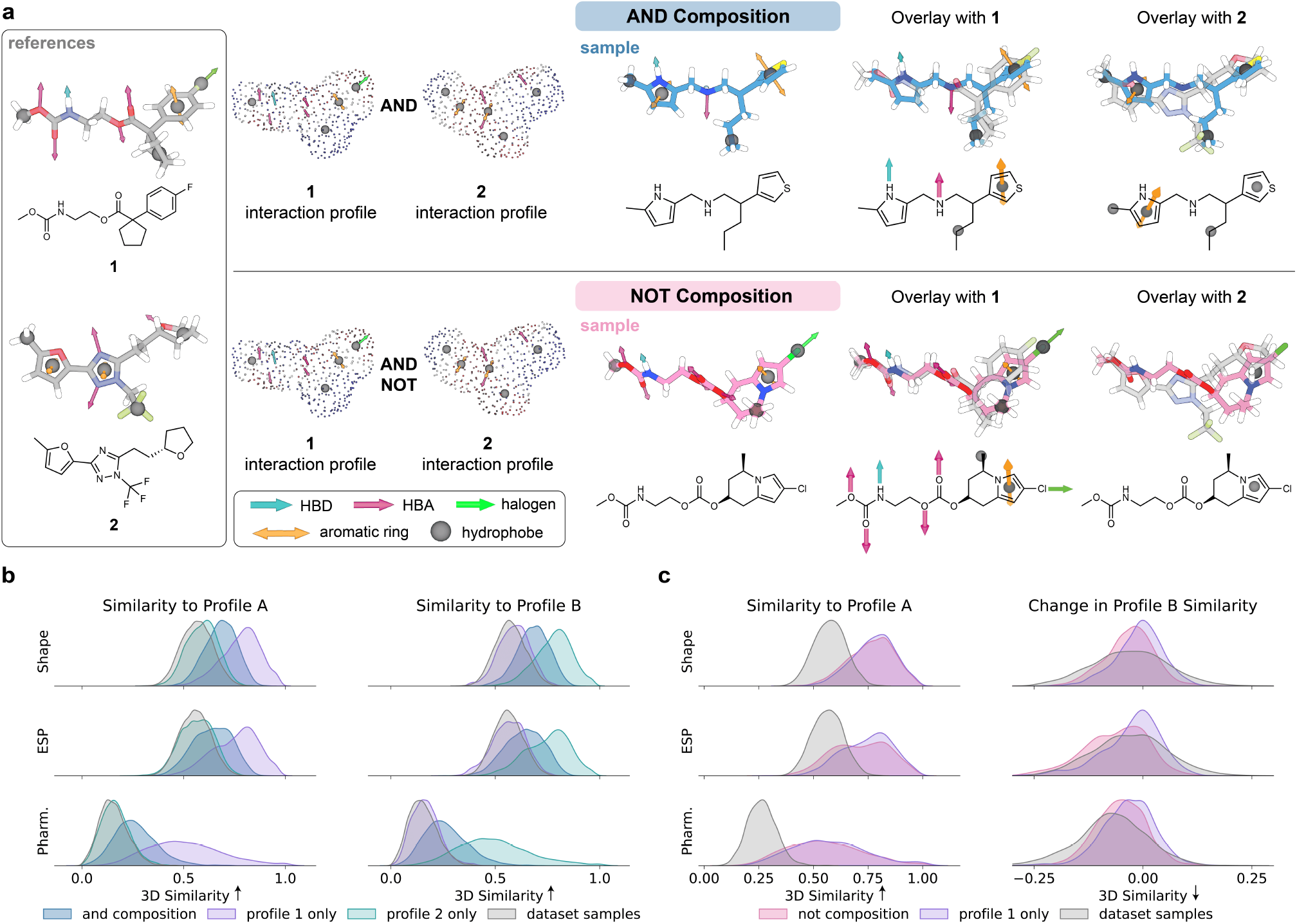
**a**, Representative schematic illustrating composition over a pair of reference profiles (gray). AND composition (blue) generates a sample compatible with both profiles (top overlay with references) while NOT composition (pink) generate a sample compatible only with the first profile (bottom overlay with references). **b**, AND composition (blue) improves the joint 3D interaction similarity to randomly paired reference profiles. **c**, NOT composition (pink) decreases similarity to the negation profile (profile 2) relative to baseline similarity while preserving similarity to its first reference profile. We note that negating random pairs is a relatively hard task, even after filtering for a minimum baseline similarity, as the input molecules’ interactions are already dissimilar from each other.

We perform a quantitative evaluation of AND and NOT composition across 100 unique random pairs of MOSES test set molecules, measuring 3D interaction similarity (shape, electrostatics, pharmacophores) to each profile. For NOT composition, we define the second profile (Profile B) of each pair as the negation profile. Composition is compared against two baselines: (1) random dataset samples as a soft lower bound of interaction similarity between two molecules, and (2) inpainting a single profile as a soft upper bound on ShEPhERD–2’s ability to recapitulate that interaction profile.

For AND composition, we observe the expected tradeoff between the interaction similarity to either pair profile, sampling a distribution intermediate to the two single-profile inpainting distributions (Fig. 4b). Inpainting solely one profile in the pair leads to close to random similarity to its paired (opposing) profile (mean Cliff’s *δ* = +0.078). Conversely, AND composition molecules have substantially enriched interaction similarity over random samples for both profiles (mean Cliff’s *δ*s = +0.588, +0.578). All comparisons are significant at *p <* 10^−4^ (Mann-Whitney; Suppl. Table S12, S13). Unlike AND, NOT composition treats the two profiles asymmetrically (Fig. 4c).

We further analyze interaction similarity for interaction profile pairs with above 0.3 baseline pharmacophore similarity (n = 24/100 pairs; Suppl. Fig. S5). This is because NOT composition presupposes shared features to negate, and therefore the possible reduction in Profile B similarity scales with a pair’s baseline similarity. Among such pairs, NOT composition’s first profile similarity is comparable to inpainting the first profile alone (mean Cliff’s *δ* = −0.089), while similarity to the negation profile is substantially reduced (mean Cliff’s *δ* = −0.247). This dissimilarity approaches or exceeds the approximate best-case lower bound set by a random distribution of samples (mean Cliff’s *δ* = −0.022) while inpainting only the first profile enriches interaction similarity over random for the negation profile (mean Cliff’s *δ* = +0.178). All comparisons between NOT composition and inpainting are significant at *p <* 10^−3^ (Mann-Whitney; Suppl. Table S14). We note that both AND and NOT composition sample molecules with modestly greater complexity and strain than single-profile generation; this is consistent with the added constraint of satisfying a composite interaction distribution, while still remaining within acceptable drug-like property ranges (Suppl. Fig. S6, S7).

### 2.5 Case studies

To demonstrate the practical utility of ShEPhERD–2 for small molecule drug design, we consider three in silico case studies. First, we design bioisosteric merges of EV-D68 3C protease fragments through an interaction-aware optimization loop. Second, we compose interactions to design dual-targeting PARP1/Tubulin inhibitors and selective BRAF V600E inhibitors. Finally, we perform modality hopping from macrocyclic peptides to small molecules that mimic their key binding interactions.

#### 2.5.1 Bioisosteric fragment merging through interaction-aware optimization

A core challenge in fragment-based drug discovery is how to grow or link fragment-sized protein-bound hits into a single coherent molecule with sufficient affinity. Numerous generative methods seek to address this by designing linkers or merges between fragment structures [48, 59, 60]. While these approaches typically preserve the fragment chemotypes, *bioisosteric fragment merging* relaxes this constraint and searches an expanded space of viable ligand hypotheses that merge the binding interactions underlying fragment affinity [36, 61]. This expanded ligand space admits scaffolds that may aggregate fragment interactions better than any linker combination between originating fragments. With ShEPhERD–2, this expanded space can be probed generatively and biased toward predicted high-affinity bioisosteric fragment merges through an interaction-aware optimization loop.

Given a set of fragment interaction profiles, we use ShEPhERD–2 to propose new molecules, score these molecules by predicted binding affinity, and seed the next iteration with the interactions of the top-scoring molecules.

Concretely, this is implemented as a genetic algorithm (GA), mutating and crossing over parent fragment interaction profiles (Suppl. Fig. S8). At each iteration, *mutation* is implemented as ShEPhERD–2 inpainting, where diverse sampled structures approximately satisfy input interactions, enabling the chemical drift expected of a mutation operator. *Crossover* is realized as AND composition over two input interaction profiles, sampling structures that attempt to satisfy both sets simultaneously. Binding affinity is evaluated by computational surrogates such as Vina docking or Boltz-2 cofolding and the predicted pose from these surrogates is used to extract interactions from top-scoring structures for subsequent iterations.

We demonstrate this interaction-aware optimization using a set of interaction profiles extracted from 13 fragments experimentally bound to EV-D68 3C protease [62], an antiviral drug target. This fragment screen was benchmarked in the original bioisosteric fragment merging work that searched a database using 2D pharmacophore fingerprints [61], enabling a direct comparison with ShEPhERD–2. We run three independent interaction-aware optimization runs (Suppl. Table S18) using docking as a binding affinity surrogate. Fig. 5a depicts the top-scoring molecule across each run. We split interaction optimization into two phases. In the exploration phase (Fig. 5a, left), the GA selects parents and evolves generated samples along the Pareto front of ligand efficiency and docking score. By construction, this enables both larger, high-affinity molecules and smaller, tightly binding fragments to persist as parents for interaction mutation or crossover operators. After 10 generations, the top 50 molecules by docking score seed the exploitation phase (Fig. 5a, right), where only docking score is optimized for a subsequent 20 generations.

**Fig. 5:**
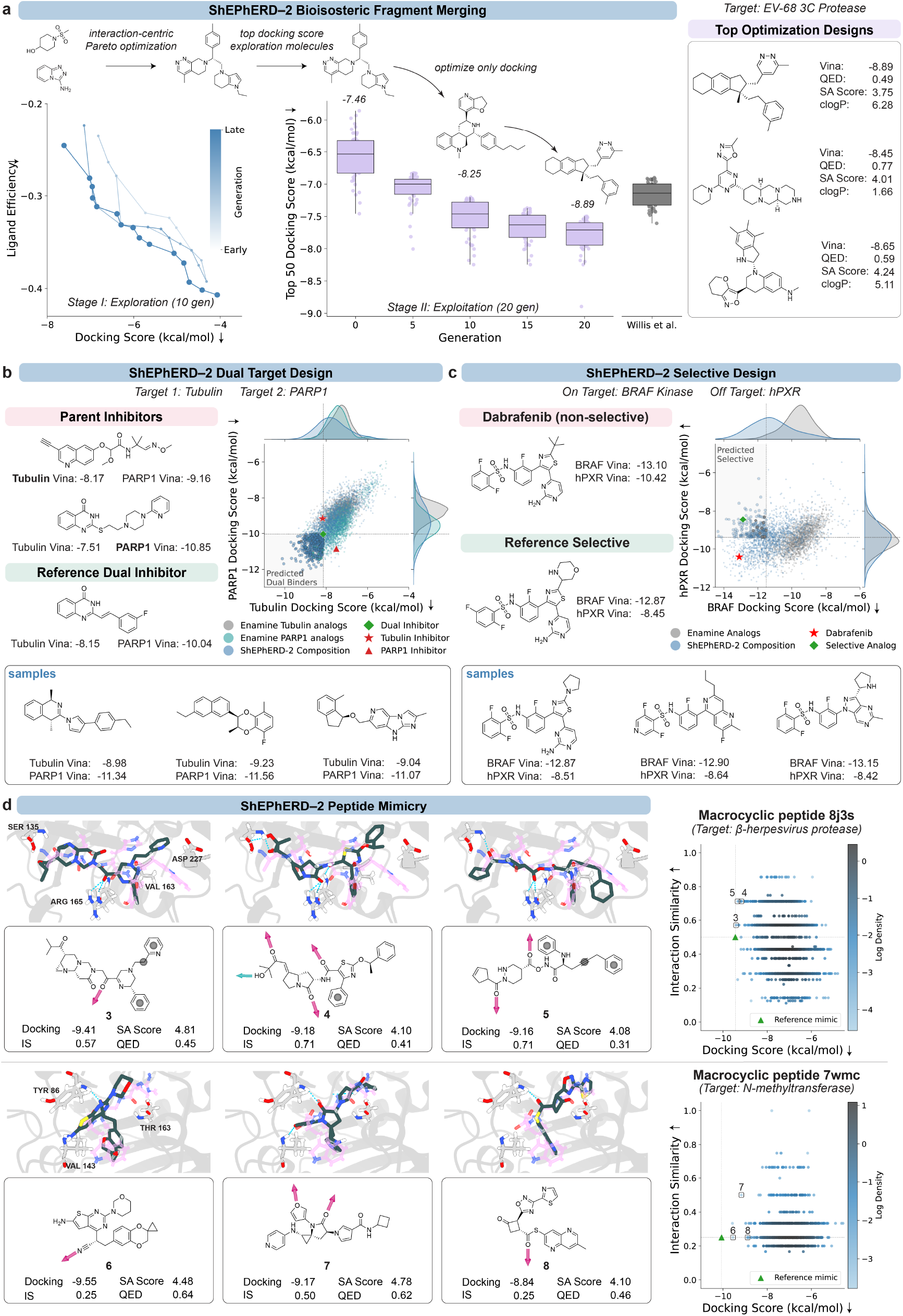
**a**, Bioisosteric fragment merging through two-stage interaction-aware optimization. Representative single run shown where stage I (left) (population = 50, generation = 10) uses a Pareto NSGA-II genetic algorithm to optimize ligand-efficient and low docking score samples seeded from EV-D68 3C fragment interactions. Interactions derived from the top 50 stage I molecules serve as gen 0 for stage II (middle) (population = 50, gen = 20), which only optimizes for docking score. Top samples across n = 3 runs shown (right). **b**, Dual target binder generation through AND composition of Tubulin and PARP1 parent inhibitor profiles. Marginal distributions for docking score to either target shown top and right for 2000 samples of AND composition and Enamine analog molecules. Putative dual binder distribution shown in the highlighted region of the plot, with representative top dual binder samples shown (bottom) compared to reference samples (top). **c**, Selective binder generation through NOT composition of dabrafenib on- and off-target bound poses. Marginal distributions shown for 2000 samples of NOT composition and Enamine dabrafenib analogs. Putative selective binder distribution shown in the highlighted region of the plot, with representative top selective binder samples shown (bottom) compared to reference samples (top). **d**, Small molecule mimetics of macrocyclic peptide interactions. Peptide interaction similarity and docking scores to peptide targets human cytomegalovirus protease and nicotinamide N-methyltransferase are shown (right). We overlay top representative sample poses (green) with the interface region of the parent peptide (pink), with key binding residues highlighted. Recovered peptide interactions detected by ProLIF are schematically highlighted in 2D molecular representations.

Interaction-aware optimization results in structures with substantially lower docking scores than the input fragments and the original bioisosteric fragment database search baseline, with optimization achieving a top-10 mean of −8.16 kcal/mol compared to the baseline top-10 mean of −7.45 kcal/mol (Cohen’s d = −4.076; Suppl. Table S18). We acknowledge, however, a tradeoff with respect to molecular properties, as database-enumerated compounds are smaller and lower in complexity than our generated compounds, though notably ShEPhERD–2 still retains comparable ligand-efficiency (Suppl. Fig. S9). Similar results arise with different choices of binding affinity surrogate such as Boltz-2 co-folding, where ShEPhERD–2 reaches a top-10 mean of −0.48 log *µ*M compared to the baseline’s 0.50 log *µ*M (Suppl. Table S19). Furthermore, interaction-aware optimization can also improve interactions seeded from prior generated samples that satisfy a manually curated aggregate fragment interaction profile (Suppl. Table S20). We refer to this as *refinement* as mutation is replaced with ShEPhERD–2 pharmacophore prioritization, reflecting local exploration of optimal interaction-satisfying structures.

#### 2.5.2 Dual target and selectivity design

Next, we consider how ShEPhERD–2 can help engineer polypharmacology and selectivity, two important levers in drug design for therapeutic efficacy and safety. In the structure-based setting, a central task in achieving polypharmacology is dual-target inhibition, while a corresponding task for selectivity is reducing a ligand’s binding to an off-target. We consider both through a ligand-centric interaction view, treating dual-target inhibition as AND composition between two parent inhibitors, and selective design as NOT composition between an on- and off-target ligand-bound pose.

We first demonstrate dual target inhibition by generating putative dual tubulin/-PARP1 inhibitors, a promising combination for cancer treatment given the synergistic effect of co-targeting tubulin and PARP1. Prior work discovered a validated dual inhibitor using a pharmacophore screen from the tubulin inhibitor 9N5 (PDB: 5O7A) and the PARP1 inhibitor MC2050 (PDB: 6XVW) [63]. We extract these same interactions from co-crystal structures and apply ShEPhERD–2’s AND composition to the pair of interaction profiles to produce 2000 molecules that attempt to satisfy both inhibitors’ interactions simultaneously. Each output is relaxed with GFN2-xTB, and RDKit-valid molecules are docked to both targets. Fig. 5b shows this joint docking distribution compared to a baseline of the 2000 nearest-neighbor structural analogs of each parent inhibitor in the Enamine REAL Database (July 2024 version, ~10 billion compounds) [64]. Compared with this “analog-by-catalog” strategy and the reference dual inhibitor from the pharmacophore screen, ShEPhERD–2 samples exhibit improved docking scores against both targets. The recovery of parent inhibitor interactions, measured by ProLIF interaction similarity, is comparable to or improved over the reference dual inhibitor for Tubulin, and closely approaches, without fully matching, the reference dual inhibitor for PARP1—unsurprising given that the dual inhibitor inherits the MC2050 substructure directly (Suppl. Fig. S10). Top dual binder samples from ShEPhERD–2 (Fig. 5b) visually show substantial deviation from their parent chemotypes, suggesting ShEPhERD–2’s broader exploration of chemical space may drive improved in silico dual affinity metrics. We acknowledge that docking is a poor surrogate of affinity and follow-up experimental validation would be necessary to claim improved dual binder activity.

We next consider NOT composition’s application to selectivity design via steering molecules away from off-target interactions. We test this with dabrafenib, an FDA-approved BRAF V600E mutant kinase inhibitor with hPXR off-target binding [65]. hPXR activation is associated with drug-drug interactions, cancer therapy resistance, and toxic metabolites, and is thus a deleterious off-target [66]. A prior medicinal chemistry campaign successfully identified selective analogs of dabrafenib, providing a retrospective benchmark for ShEPhERD–2 [67].

Bound poses of dabrafenib to BRAF V600E kinase (PDB: 4XV2) and hPXR (PDB: 6HJ2) were used to define interaction profiles for NOT composition, with the latter used as the negation profile. Since negating a compound with itself is ill-defined, we subselected the interactions with differential favorability for the off-target using Vina (Suppl. Sec. S1.12). We then applied NOT composition to this pair to generate structures that satisfy on-target interactions while favoring off-target-incompatible chemotypes. In total, 2000 samples were generated, and valid samples post-relaxation were docked to both proteins. As shown in Fig. 5c, ShEPhERD–2 samples are enriched for similar on-target docking scores alongside significantly worse off-target docking scores compared to the 2000 nearest analogs of dabrafenib in Enamine REAL, most of which do not contain the pyrimidine motif important for BRAF binding. While docking may not reliably score off-target binders [68], these results demonstrate that NOT composition can steer molecules toward interactions that disrupt previously predicted off-target binding modes. Notably, ShEPhERD–2 NOT composition recovers the known selective analog’s design strategy of replacing dabrafenib’s tert-butyl with a bulky polar substituent. Visual inspection reveals that 13 of the top 50 most-selective samples (by differential docking score to BRAF) exhibit this replacement (Suppl. Fig. S11), with a representative example of a pyrrolidine rather than the analog’s morpholine shown in Fig. 5c. In silico selective redesigns of dabrafenib can also be generated with differing levels of fidelity to the known active scaffold using scaffold conditioning. NOT composition with conservation of a known active substructure shared across BRAF inhibitors was able to achieve similar selective enrichment against BRAF and hPXR (Suppl. Fig. S12).

#### 2.5.3 Modality hopping

The universality of using molecular interactions as the basis for ShEPhERD–2’s conditional molecular generation enables agnosticism about the modality from which these interactions are derived. This allows us to consider a class of problems where a non-small-molecule compound (e.g., macrocycle, peptide, protein, nucleic acid) has pharmacological utility from its interactions but liabilities from its structure. With ShEPhERD–2, we can “project” such compounds to drug-like small molecules that mimic their interactions. We refer to this general procedure as *modality hopping*.

One such common modality hop in drug discovery is peptidomimicry or peptide-to-small-molecule conversion [69–72]. Peptidomimicry campaigns aim to overcome the disadvantages of peptides, such as poor oral bioavailability, cell permeability, or patient adherence, via the discovery of a small molecule mimic of peptide interactions. We test ShEPhERD–2’s ability to modality hop from peptide to small molecule in silico using two macrocyclic peptides targeting human cytomegalovirus protease (HCMV-Pro) and nicotinamide N-methyltransferase (NNMT), respectively [73, 74]. Each of these peptides has a reference small-molecule mimic of its core binding interactions as a basis for comparison (Suppl. Fig. S13). For HCMV-Pro, these are hydrogen bond acceptors from Ser135 and Arg165, hydrogen bond donors to Ser135, and hydrophobic interactions with Val163, Ser135, and Asp227. For NNMT, they include hydrogen bond donors to Thr163, hydrogen bond acceptors from Tyr86 and Val143, and hydrophobic interactions with Val143.

These same peptide interactions were used to define the input interaction profiles for ShEPhERD–2. For each peptide, we generated 2000 samples using pharmacophore conditioning across a 20-atom range, relaxed with GFN2-xTB to filter invalid or unstable compounds, and docked them to the target protein using Vina. Fig. 5d shows the samples’ docking scores and ProLIF interaction similarities (IS) of the docked pose to these input interactions. Compared to the reference mimic, ShEPhERD–2 was able to sample peptide mimics with comparable docking scores and improved interaction similarity without significantly lower QED or SA scores (Suppl. Table S21). Visual inspection of top-generated samples by docking score and IS in Fig. 5d reveals diverse structural hypotheses for mimicry, providing multiple candidate starting points for medicinal chemistry optimization. Interestingly, molecules with the largest IS often yielded worse docking scores than lower-IS samples. Many of the generated molecules that achieve high interaction similarity do so through linear peptide-like structures whose conformational entropy is penalized by Vina’s scoring function (Suppl. Fig. S14), suggesting that perfect interaction similarity may not necessarily constitute the best mimics for downstream peptidomimicry campaigns.

## 3 Discussion

Three-dimensional intermolecular interactions between molecules and their environment govern molecular function and are central to molecular design. Despite this, many models for molecular generation consider intermolecular interactions only implicitly, despite their explicit treatment during subsequent steps of evaluation or visualization, including by expert medicinal chemists. To better align generative design with the field’s rich knowledge of interactions that drive target engagement, we introduced ShEPhERD–2, a flexible generative framework that considers contributions of sterics, electrostatics, and non-covalent interactions *during* generation. This 3D generative model advances interaction-aware ligand-based molecular generation while approaching the performance of structure-based models despite representing the target pocket only through its bound ligand interactions. By operating directly on interactions rather than specific chemical structures, ShEPhERD–2 provides a common framework for diverse design tasks spanning interaction-based optimization, structure-informed design, and modality hopping. Innovations to the model enable a high degree of controllability for prioritizing pharmacophore subsets, generating molecules with conserved substructures, and composing multiple interaction profiles. Together, these capabilities support a broader view of interaction profiles as a universal design language that is independent of molecular scaffold, target representation, or chemical modality.

Several opportunities remain to further extend this framework. First, our prospective case studies rely on docking and interaction analyses as a poor proxy for binding affinity, whereas experimental validation will ultimately be required to confirm the intended function. Second, interaction profiles currently originate from ligands with known activity. Methods that infer interaction constraints directly from protein binding pockets [47, 55, 75–80] could enable interaction-conditioned *de novo* design to explore beyond known recognition patterns. Finally, although interaction-conditioned generation enables exploration beyond enumerated chemical libraries, synthetic accessibility remains an important consideration. Post-hoc shape-conditioned database search [81] or generative projection into a theoretically synthetically accessible space [82–84] can help ameliorate this limitation but risks relaxing away the precisely designed interactions. As Rekesh et al. [85] has begun to explore, incorporating synthetic feasibility directly into an interaction-aware generative process, rather than as a post hoc projection, represents an important direction for future work.

More broadly, we believe this work demonstrates how interaction profiles provide a transferable representation of molecular function that is largely independent of the underlying molecular system. The interactions that govern protein–ligand recognition arise from the same physical principles that underlie molecular recognition across chemistry, including supramolecular host–guest complexes and catalyst–substrate interactions. Future extensions that incorporate additional determinants of molecular function may enable interaction-conditioned generative models to serve as a common framework for molecular design across diverse chemical domains.

## 4 Methods

### 4.1 Representations

ShEPhERD–2 operates on a joint representation ***X*** of 3D molecular structure ***x***_1_, shape ***x***_2_, electrostatic potential ***x***_3_, and pharmacophoric features ***x***_4_ as introduced by ShEPhERD [36]. The molecular structure is represented as a 3D molecular graph containing atom types, formal charges, covalent bond orders, and atomic coordinates, including explicit hydrogens. Molecular shape is represented as points sampled on the solvent-accessible surface, and electrostatic potential (ESP) is represented by the Coulombic potential at each surface point using GFN2-xTB-computed atomic partial charges. Pharmacophoric features are extracted from SMARTS patterns compiled from established pharmacophore definitions [86–89]. These include directional features, such as hydrogen bond donors and acceptors, aromatic rings, and halogen bonds, and directionless features, such as hydrophobes, anions, cations, and zinc binders. Each pharmacophoric feature is represented by its type, position, and, when applicable, a unit vector for its directionality. Exact definitions are found in Suppl. Sec. S1.1.

### 4.2 Generative diffusion framework

ShEPhERD–2 models the joint distribution of 3D molecular structures and interaction profiles using a variance-exploding (VE) diffusion process with EDM parameterization [90]. The model is trained to recover the clean joint representations from Gaussian-corrupted examples and, at inference, generate new samples by iterating this denoising operation over decreasing noise levels. Notably, all variables are treated as continuous, with discrete variables represented as continuous one-hot encodings. We summarize the generative procedures below and reserve detailed explanations for Suppl. Sec. S1.2. Given a clean joint representation ***X***^(0)^, the forward noising process independently adds Gaussian noise to each variable at a uniform noise level *σ* resulting in a noisy representation ***X***^(*σ*)^ = ***X***^(0)^ + *σ* ***ϵ***, where ***ϵ*** ~ *N* (**0, *I***). At sufficiently large *σ*, the resulting distribution is approximately an isotropic Gaussian, which serves as the sampling prior. Atomic-coordinate noise is projected to remove its center of mass (COM) to preserve translational invariance [91]; the coordinates of the other modalities are not independently recentered so that their spatial relationship to the molecular structure is retained. The denoiser *D*_*θ*_ is trained with a weighted L2 loss to recover the clean representation from its noisy counterpart: 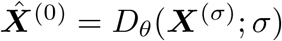.

At inference time, we use a non-Markovian predict–renoise procedure [92, 93]. Starting from 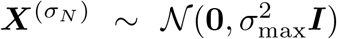, we iteratively proceed through decreasing noise levels to *σ*_min_. At each denoising step, ShEPhERD–2 predicts the clean, joint representation and then adds fresh Gaussian noise at the next noise level:

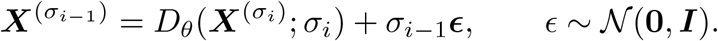

In practice, we generate molecule structures conditioned on interaction profiles through complementary strategies. These include inpainting, fixed pharmacophore and atom conditioning, and composition.

#### Interaction inpainting

The representation is partitioned into a “known” interaction profile context, ***X***_*K*_, and the structure to be generated, ***X***_*G*_. Initially, we simulate and store a forward noising trajectory of the conditioning context 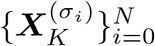. During predict–renoise sampling, we replace 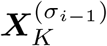 with the corresponding simulated context and retain the model’s re-noised prediction 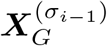 such that 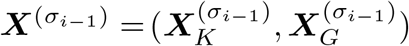.

#### Fixed pharmacophore and substructure conditioning

To enable conditioning on fixed pharmacophores or substructures ***X***_*F*_, we randomly select pharmacophore types, positions, and directions, or atom types, formal charges, and positions to remain unnoised during training. These fixed nodes are masked from the loss, and optimization yields the denoiser 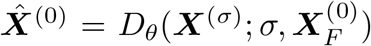. The system is centered on either the noised COM of the molecular structure or the COM of the fixed points, and the atomic coordinate noise is not projected to remove its COM. ***X***_*F*_ is fixed throughout denoising, and we optionally inpaint other known features that are not fixed.

#### Composition

For AND and NOT composition, we follow the formulation established by Liu et al. [58] and prove that it applies to our predict–renoise sampler in Suppl. Sec. S3.1.1-S3.1.2. Given 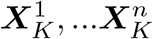 desired interaction profiles, we conditionally sample a single molecular structure that attempts to satisfy all profiles by inpainting them with the AND compositional sampler [58]:

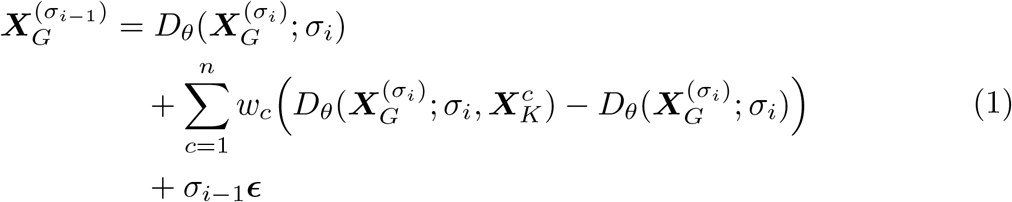

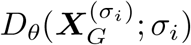 is the unconditional denoised prediction, while 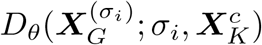 is the denoised prediction under inpainting 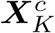. Scalar weights *w*_*c*_ control the contribution of each profile to the denoised prediction. For NOT composition, we adopt a similar formulation to favor an on-target interaction profile ***X***_*on*_ and disfavor off-target profiles 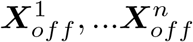. Following Liu et al. [58], we define the NOT compositional sampler:

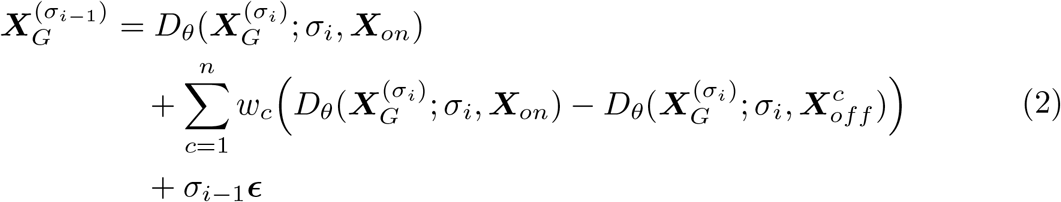

### 4.3 ShEPhERD–2 architecture

ShEPhERD–2 adopts the overall SE(3)-equivariant design of ShEPhERD, which consists of (1) separate *embedding modules* for each representation into invariant (*ℓ* = 0) and equivariant (*ℓ* = 1) features, (2) a *joint module* that operates on a cross-modal heterogeneous graph, and (3) separate *denoising modules* for each modality. Details are documented in Adams et al. [36], while key updates are described below.

We modify the ShEPhERD backbone by replacing the EquiformerV2-style SE(3)-equivariant graph transformer modules [94] with the faster and more expressive EquiformerV3 modules [95]. Following Liao et al. [95], we favor the more expressive SwiGLU nonlinear activation over SiLU, enabling a coarser grid resolution to further improve model efficiency. We reduce the expressivity of the global *ℓ* = 1 channel embedding, constructed in the joint module, prior to taking the tensor product with the noise embedding to reduce overfitting. In the setting where there is fixed conditional context ***X***_*F*_ (Suppl. Sec. 4.2), we add learnable embeddings to the *local*, invariant *ℓ* = 0 latents of the corresponding fixed nodes within the embedding module. Furthermore, we add another learnable embedding to the *global ℓ* = 0 latent embeddings in the joint module. Model parameters are documented in Suppl. Table S2.

### 4.4 ShEPhERD–2 training

We trained ShEPhERD–2 on the MOSES dataset, which contains 1.6 million neutral, drug-like molecules with up to 27 heavy atoms [38]. In particular, we use the ShEPhERD-MOSES-aq version prepared by Adams et al. [36], in which each molecule is represented by a single conformer with explicit hydrogens, locally relaxed using GFN2-xTB in implicit aqueous solvent [37]. The dataset contains H, C, N, O, F, Cl, Br, I, S, P, and Si element types. We remove the unweighted center of mass (COM) from each conformer, then compute its surface, electrostatic potential, and pharmacophore representations. Each training example is augmented by adding up to 10 dummy atoms and 5 dummy pharmacophoric features. These features are placed on the noised coordinates of randomly selected atoms and pharmacophoric features, respectively [39, 40].

We sample the noise level *σ* from a log-normal distribution, 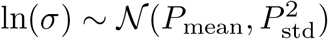 [90], and independently noise each variable with isotropic Gaussian noise, *ϵ* ~ *N* (**0**, *σ*^2^***I***). For the setting of fixed atoms and pharmacophores, random atoms and pharmacophores remain unnoised and are masked from the loss. We train ShEPhERD– 2 using a weighted L2 loss for 300k optimization steps using Adam optimizer with an effective batch size of 96. Suppl. Sec. S1.2 and S1.7 contain detailed descriptions of training.

### 4.5 Evaluations

#### 4.5.1 MOSES test set experiments

Our test set is composed of 100 neutral, drug-like molecules randomly selected from the MOSES scaffold-split test set, which contains Bemis-Murcko scaffolds not found in the training set [38]. The conformers were prepared in the same way as the training set: a single conformer relaxed with GFN2-xTB in implicit aqueous solvent. We generate with the sampling parameters described in Suppl. Sec. S1.9.

##### Evaluations

Each generated sample is evaluated through a set of conformer quality metrics after local relaxation of the extracted structures with GFN2-xTB. The energy difference between the pre- and post-relaxed structures defines the strain energy. A sample is considered chemically valid if RDKit v2026.3.1 [96] can construct a molecular graph from the relaxed XYZ coordinates. This molecular graph is used to compute auxiliary metrics such as graph similarity to the reference molecule via Morgan fingerprint (radius 3, 2048 bits, with chirality) Tanimoto similarity, synthetic accessibility (SA) score [97], quantitative drug-likeness score (QED) [98], cLogP, and the fraction of sp3 carbons 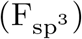.

We extract the interaction profile from the relaxed conformation and compute 3D interaction similarity to the reference interaction profile for each interaction modality, *i* ∈ {2, 3, 4}. Specifically, we use the Tanimoto similarity of pairwise, first-order Gaussian overlaps between the point cloud representations of an interaction modality for the sampled structure 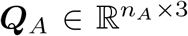 with *n*_*A*_ points and the reference 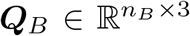 with *n*_*B*_ points: sim_*i*_(***Q***_*A*_, ***Q***_*B*_) ∈ [0, 1] [13, 14, 36, 99, 100]. Since 3D similarity depends on the relative orientations of the conformers, we report the optimal similarity, 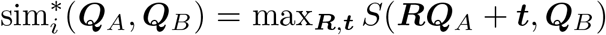 obtained by optimizing over rotations ***R*** ∈ *SO*(3) and translations ***t*** ∈ *T* (3). We perform *local* alignment by optimizing from the sampled (post-relaxation) pose for up to 200 steps with the Adam optimizer. When specified, we may perform *global* optimization, which simultaneously aligns 50 randomly seeded poses and reports the highest score. To compute the 3D similarity of a *subset* of an interaction profile (e.g., high priority or low priority pharmacophore), we utilize Tversky similarity computed after alignment to the *full* profile. The exact scoring function used for each modality is detailed in Suppl. Sec. S1.8.

##### Interaction-conditioned generation

For benchmarking interaction-conditioned analoging and hit expansion, we generate 20 samples for each test molecule by inpainting its associated interaction profile. Unless otherwise specified, we initialize the number of atoms and pharmacophores to be the same as the reference: 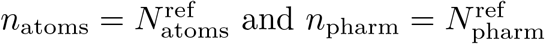.

##### Pharmacophore conditioning

To evaluate ShEPhERD–2’s ability to condition on fixed pharmacophore features, we randomly select 30–50% of the pharmacophore features of each reference to be high-priority (HP) and supply their unnoised pharmacophore types, positions, and directions to the model throughout generation. We consider three generative settings: “HP only,” where we fix the HP pharmacophore features throughout generation and ignore the rest; “prioritization,” where we fix the HP pharmacophore features and inpaint the rest; and “all fixed,” where we fix the entire reference pharmacophore. In all cases, we also inpaint surface and ESP. After GFN2-xTB relaxation of the samples, we score the full 3D pharmacophore similarity after local alignment to the reference molecule’s pharmacophore. Using this relaxed and aligned pose, we compute the Tversky similarity for both HP and low priority (LP) subsets. Note that Tanimoto and Tversky similarities are not directly comparable.

##### Substructure conditioning

To test ShEPhERD–2’s ability to condition on fixed atoms or substructures, we consider two settings. First, we randomly select heteroatoms including neighboring hydrogens, or if the heteroatom is terminal, its heavy-atom neighbor. Second, we use BRICS fragmentation [49] and randomly select a single fragment containing at least 10 atoms. For these selected atoms, we supply unnoised atom types, formal charges, and positions throughout the diffusion trajectory. Alongside fixed scaffold conditioning, we inpaint the full interaction profile.

##### Composition

We generate 100 random, unique pairs from the MOSES test set and generate 80 samples per pair across 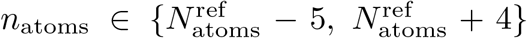 and 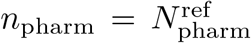. For both forms of composition, 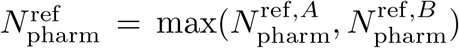, while we define 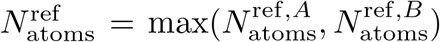 for AND composition and 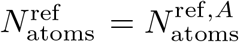 for NOT composition. A shared reference frame is calculated between paired molecules through global alignment of ESP similarity. In all cases, we inpaint using pharmacophores, surface, and ESP with equal condition weighting of 0.5 during composition (Eq. 1, 2). After sample relaxation with GFN2-xTB, we score 3D interaction similarity following local alignment to each paired reference, while negation profiles (profile 2 in NOT composition) are scored following global alignment.

##### Uncertainty quantification

For each metric and setting, we perform two-sided Mann-Whitney U tests for pairwise distributions, pooling generated molecules across the test set. *p*-values are Holm-adjusted for each metric. Effect sizes are reported as Cliff’s *δ* = *P* (*A > B*) − *P* (*A < B*). Validity is reported as the mean per-reference validity rate, with 95% confidence intervals computed using Student’s *t*-distribution.

#### 4.5.2 MolGenBench hit-to-lead benchmark

For each of the 600 reference molecules in the MolGenBench hit-to-lead benchmark [51], we condition generation on the interaction profile extracted from the bound poses. Because ShEPhERD–2 was trained only on neutral molecules and some reference molecules are charged, we reduce train–test mismatch of the electrostatic potential (ESP) profile by uniformly removing the formal charge, 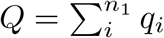, across all atoms The Coulombic potential is then recomputed from 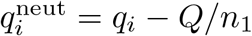.

We increase sampling coverage by initializing samples with additional atoms and pharmacophoric features relative to the reference molecule. For each reference, we sample 10 batches of 20 molecules using the settings [(0, 0), (1, 0), (2, 0), (2, 1), (2, 2), (3, 0), (3, 1), (3, 2), (4, 1), (4, 2)], where each tuple specifies the number of additional atoms and pharmacophores, respectively.

In the pocket-conditioned setting, we additionally extract pharmacophore features associated with ProLIF-detected non-covalent interactions between the ligand and the pocket [101]. These high-priority features are fixed during generation, while the remaining pharmacophore features, surface, and ESP are inpainted, constituting pharmacophore prioritization. As an ablation, we fix interacting ligand atoms that form the ProLIF-detected non-covalent interaction instead of pharmacophore prioritization.

Alongside existing MolGenBench evaluation metrics, we introduce an “Interaction Diversity” metric. This captures the mean chemical diversity of sampled substructures at ProLIF-identified interactions. For each interaction present in *>*10% of the reference actives, the ligand atoms involved are used to define a surrounding substructure within a 3-bond radius, from which a Morgan fingerprint (radius 2, 2048 bits, without chirality) is computed. Interaction diversity is calculated as one minus the mean pairwise Tanimoto similarity across generated molecules, and then averaged across interactions for each target protein.

We evaluated the baseline models using their standard, interaction-conditioned settings, except for PhoreGen, where we modified the sampling procedure to help the model condition on pharmacophore representations larger than those in its training set in the ligand-based setting. Further experimental details on the protocols for each baseline and the definitions of MolGenBench metrics are reported in Suppl. Sec. S1.10.

#### 4.5.3 Case studies

##### Docking and co-folding evaluations

Case study target receptors were prepped using UCSF ChimeraX Dock Prep [102] from their reference inhibitor-bound structures. Exact choices of reference are delineated in each case study. Docking was performed with AutoDock Vina 1.2.6 [103] using a 20 Å × 20 Å × 20 Å grid centered on the reference inhibitor’s center of mass and an exhaustiveness of 32. We follow the docking pipeline outlined in the original ShEPhERD work [36], where protonation states are preserved from samples while conformers are re-enumerated from SMILES. For co-folding evaluations, we employed Boltz-2 [104] with 3 recycling steps, 1 diffusion sample, 200 sampling steps, multiple-sequence alignment, and steering potentials enabled. Protein sequences for Boltz-2 were extracted directly from the prepped structures after removal of histidine tags.

##### Bioisosteric fragment merging

To search for optimal bioisosteric fragment merges, we implement a genetic algorithm (GA) that operates on interaction profiles. Each individual ℐ = (***X***, *s*) couples a structure ***x***_1_ to its interaction profile (***x***_2_, ***x***_3_, ***x***_4_) with predicted binding affinity fitness *s*. At each iteration, individuals ℐ selected on the basis of *s* undergo mutation and crossover over their interaction profiles to generate new individuals ℐ^′^. We outline further details of the genetic algorithm in Suppl. Sec. S1.11.

Our initial fragment population 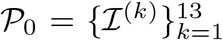 comprises 13 fragments from an EV-D68 3C fragment screen from Diamond Light Source [105] selected as seeds in Adams et al. [36], with interaction profiles extracted from the native pose. We set the per-generation GA population *P* = 50 with individuals added to *P* at generation *g* if the generated structure ***x***_1_ satisfies Lipinski’s Rule of Five, is free of PAINS and Brenk substructure alerts, has fewer than 10 rotatable bonds, and has an SA Score less than 4.5. Binding fitness *s* is scored for each individual based on the EV-D68 3C docking score to the apo structure (PDB: 8CNX) or the Boltz-2 predicted binding affinity using the EV-D68 3C sequence.

We split bioisosteric fragment merging into an exploration and exploitation phase with different GA parameters (Suppl. Sec. S1.11.3). In the exploration phase, we (1) enable interaction crossover to generate ℐ^′^ with 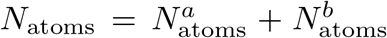 if both *a* and *b* are fragments of fewer than 25 heavy atoms, (2) increase the probability of interaction crossover to 0.8, (3) merge with a unique original fragment with probability *p*_*o*_, and (4) select parent individuals ℐ for mutation and crossover at each *g* based on non-dominated sorting (NSGA-II) [106] over (*s, s*_ligand−efficiency_). We find that multi-objective Pareto optimization (Suppl. Sec. S1.11.2) with ligand efficiency is necessary to prevent premature convergence to larger input fragment merges. For Boltz-2, we transform the output log *µ*M units into kcal/mol since ligand efficiency rankings are not invariant to affine transformations. Following 10 generations of exploration, we extract the top 50 ℐ by *s* to serve as *P*_0_ for the exploitation phase, optimizing solely *s* with ℐ = 50 and *g* = 20 and obeying the default atom count rules of Suppl. Sec. S1.11.1. In either phase, selection is by tournament selection of size 8 from the cumulative pool of the top 50 ℐ, and interaction profiles for subsequent *g* are extracted from the predicted poses output by docking or cofolding. We note that exploration and exploitation parameters are adjustable and may need to be tuned in a target-dependent fashion.

##### Dual target and selectivity design

To evaluate AND composition for dual target inhibition, we generated 2000 samples from the ESP globally aligned interaction profiles of a Tubulin inhibitor 9N5 (PDB: 5O7A) and a PARP1 inhibitor MC2050 (PDB: 6XVW). We define 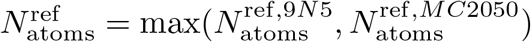 and sample uniformly 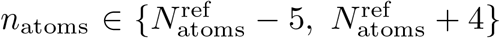 and 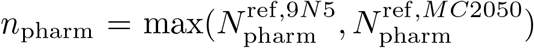 with a composition weight of 0.5 for each profile. We relaxed all samples with GFN2-xTB before assigning bond orders with RDKit and filtering chemically invalid samples. Docking to either protein target was conducted using the reference inhibitor-bound structures. For baseline comparisons, we retrieved reference inhibitor analogs through the Enamine REAL Database (July 2024 version, ~10 billion compounds) [64] by ECFP4 Tanimoto similarity. The 2000 nearest analogs were ranked by Tanimoto similarity and filtered to ensure they had no more heavy atoms than the maximum among ShEPhERD–2 samples.

For NOT composition, a similar procedure was performed for selectivity design. We generated 2000 samples from the ESP globally aligned interaction profiles of dabrafenib’s on-target BRAF V600E kinase pose (PDB: 4XV2) and off-target hPXR pose (PDB: 6HJ2), with the associated receptor structures used in later docking evaluations. The off-target pose interaction profile was sub-selected using heavy atoms that were differentially favorable for the off-target using Vina’s scoring function (Suppl. Sec. S1.12). We define 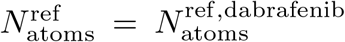 and sample uniformly 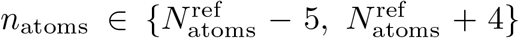 and 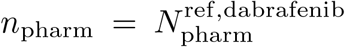 with composition weights 0.5, *™*0.5. We retrieved the 2000 nearest Enamine analogs to dabrafenib in the same manner as for dual-target inhibition.

##### Modality hopping

Macrocyclic peptide interaction profiles were determined from sub-selected interface interactions outlined in prior small molecule mimicry work [73, 74]. These interactions were HBA to Ser135 and Arg165, HBD to Ser135, and HYD with Val163, Ser135, and Asp227 for human cytomegalovirus protease (PDB: 8J3S) and HBD to Thr163, HBA to Tyr86 and Val143, and HYD with Val143 for nicotinamide N-methyltransferase (PDB: 7WMC). We include all associated pharmacophore atoms and linking backbone atoms between pharmacophores in the shape representation and add a 1.5 Å radial buffer to reduce linear shapes. We inpaint only the shape and pharmacophores and set *σ*_*max*_ to 30 to reduce amide formation and encourage ShEPhERD–2 to sample molecules that reach the spread-out peptide interactions. We set 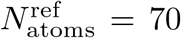 for human cytomegalovirus protease and 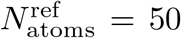 for nicotinamide N-methyltransferase and sample 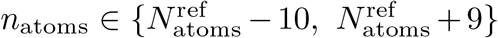. Since pharmacophores are sub-selected across the interface while shape is not, we sample *n*_pharm_ from an empirical distribution *P* (*n*_pharm_ | *n*_atoms_) constructed from the ChEMBL small molecule approved drug dataset [107]. We generate 2000 samples from each profile and dock against holo structures of the target receptors obtained from peptide-bound reference structures.

## Acknowledgments

The authors thank Ne Dassanayake and Shrey Goel for contributions to improving the formulation and code of ShEPhERD–2. We also thank Ross Irwin for insightful discussions on practical molecular generation. Finally, we acknowledge Dr. Jenna C. Fromer, Matthew Cox, and Shitong Luo for valuable insight and feedback throughout model development.

## Data availability

Model weights are on HuggingFace https://huggingface.co/kabeywar/shepherd2. Training data and model outputs used for evaluations are available on Zenodo https://doi.org/10.5281/zenodo.22675706.

## Code availability

All code is freely available under MIT license. The ShEPhERD– 2 source code and scripts used for generation are found on GitHub at https://github.com/coleygroup/shepherd2. Evaluations were facilitated by the ShEPhERD-score package v1.4 found at https://github.com/coleygroup/shepherd-score.

## Author contribution

All authors contributed to the conceptualization of this project. K.A.A. developed the ShEPhERD–2 model and led model evaluations. K.W. developed the composition algorithms and led the case studies. All authors contributed to the writing of this paper.

## Footnotes

1 **Sh**ape, **E**lectrostatics, and **Ph**armacophore **E**xplicit **R**epresentation **D**iffusion

