## Supplemental Information for "Interaction Profiles as a Universal Language for Generative Molecular Design with ShEPhERD–2"

### S1 Supplemental Methods

#### S1.1 Representations

ShEPhERD–2 operates on a joint representation of 3D molecular structure $\boldsymbol{x}_1$, solvent-accessible surface $\boldsymbol{x}_2$, Coulombic potential $\boldsymbol{x}_3$, and directionless (e.g., hydrophobic, ionic) and directional (e.g., hydrogen bond acceptors/donors, $\pi$ interactions) pharmacophoric features $\boldsymbol{x}_4$. Collectively, we define these representations as $\boldsymbol{X} = (\boldsymbol{x}_1, \boldsymbol{x}_2, \boldsymbol{x}_3, \boldsymbol{x}_4)$.

##### S1.1.1 Molecular structure

We define the 3D molecular structure (including explicit hydrogens) of an organic molecule with $n_1$ atoms as a molecular graph $\boldsymbol{x}_1 = (\boldsymbol{a}, \boldsymbol{f}, \boldsymbol{C}, \boldsymbol{B})$ with one-hot encodings for categorical features: atom types $\boldsymbol{a} \in \mathbb{R}^{n_1 \times N_a}$ with $N_a$ atom type options; formal charges $\boldsymbol{f} \in \mathbb{R}^{n_1 \times 5}$, which range from $-2$ to 2; and the covalent bond order adjacency matrix $\boldsymbol{B} \in \mathbb{R}^{n_1 \times n_1 \times 5}$, which includes no bond, single, double, triple, and aromatic bonds. $\boldsymbol{C} \in \mathbb{R}^{n_1 \times 3}$ are the continuous atomic coordinates.

##### S1.1.2 Shape

We define the shape $\boldsymbol{x}_2 = \boldsymbol{S}_2 \in \mathbb{R}^{n_2 \times 3}$ as a point cloud of $n_2$ points sampled from the solvent-accessible surface of $\boldsymbol{x}_1$. We represent the shape as the *surface* to better decouple the shape representation from the molecule's exact atomic coordinates or 2D graph. $n_2$ is fixed at 75 during training and inference for computational efficiency, but is upsampled to 400 for 3D shape similarity evaluations.

##### S1.1.3 Electrostatic potential

We define the electrostatic potential $\boldsymbol{x}_3 = (\boldsymbol{S}_2, \boldsymbol{v})$ by the Coulombic potential computed at each of the $n_2$ points sampled on the solvent-accessible surface $\boldsymbol{S}_2 \in \mathbb{R}^{n_2 \times 3}$. We compute $\boldsymbol{v}$ from the GFN2-xTB partial charges of each atom in $\boldsymbol{x}_1$.

##### S1.1.4 Pharmacophore

We extract pharmacophoric features from SMARTS patterns compiled from Koes and Camacho [1], Landrum et al. [2], Kutlushina et al. [3] with hydrophobe clustering following Berenger and Tsuda [4]. A set of $n_4$ pharmacophoric features is defined as $\boldsymbol{x}_4 = (\boldsymbol{p}, \boldsymbol{P}, \boldsymbol{V})$ where $\boldsymbol{p} \in \mathbb{R}^{n_4 \times N_p}$ is a one-hot encoding of $N_p$ pharmacophore types; $\boldsymbol{P} \in \mathbb{R}^{n_4 \times 3}$ contains the coordinates of each pharmacophore feature; and $\boldsymbol{V} \in \{\mathbb{S}^2, \boldsymbol{0}\}^{n_4}$ is the set of relative unit or zero vectors specifying the directionality of each feature, capturing directional (hydrogen bond acceptors, hydrogen bond donors, aromatic rings, and halogen bonds) and directionless (hydrophobes, anions, cations, zinc binders) pharmacophoric features. To simplify the diffusion generative process, we map the directional vectors to $\boldsymbol{V} \in \mathbb{R}^{n_4 \times 3}$.

#### S1.2 Generative diffusion framework

We model the joint distribution of molecular structures and interaction profiles, $\boldsymbol{X}$, using a variance-exploding (VE) diffusion process under the EDM parameterization [5]. The model is trained to recover clean joint representations from Gaussian-corrupted examples and, at inference, generates new samples by iterating this denoising operation over decreasing noise levels.

##### S1.2.1 Variance-exploding forward noising

Let $\boldsymbol{X}^{(0)} \sim p_0 = p_{\text{data}}$ be the clean joint representation. With some abuse of notation, the VE noising process adds isotropic Gaussian noise to the clean sample,

$$\boldsymbol{X}^{(\sigma)} = \boldsymbol{X}^{(0)} + \sigma\,\boldsymbol{\epsilon}, \qquad \boldsymbol{\epsilon} \sim \mathcal{N}(\mathbf{0}, \boldsymbol{I}), \tag{S1}$$

where $\sigma \in [\sigma_{\min}, \sigma_{\max}]$ controls the corruption level. Accordingly, the transition kernel $q_\sigma$ and the marginal density $p_\sigma$ are given by

$$q_\sigma(\boldsymbol{X}^{(\sigma)} \mid \boldsymbol{X}^{(0)}) = \mathcal{N}(\boldsymbol{X}^{(0)}, \sigma^2\boldsymbol{I}), \qquad p_\sigma = p_0 * \mathcal{N}(\mathbf{0}, \sigma^2\boldsymbol{I}). \tag{S2}$$

When $\sigma_{\max}$ is large relative to the scale of the data, we approximate $p_{\sigma_{\max}}$ by $\mathcal{N}(\mathbf{0}, \sigma_{\max}^2\boldsymbol{I})$ and use this as the sampling prior.

We sample independent noise vectors $\boldsymbol{\epsilon}_m$ for each variable $m$ defined in Sec. 4.1. Notably, we symmetrize the bond adjacency matrix $\boldsymbol{B}^{(\sigma)}$ and we remove the center of mass (COM) from the noise added to the atomic coordinate channel $\boldsymbol{C}$ for translational invariance. If $\boldsymbol{\epsilon}_C \in \mathbb{R}^{n_1 \times 3}$ is the initially sampled atomic coordinate noise, we use

$$\tilde{\boldsymbol{\epsilon}}_C = \boldsymbol{\epsilon}_C - \frac{1}{n_1}\sum_{j=1}^{n_1} \boldsymbol{\epsilon}_{C,j}. \tag{S3}$$

Only the coordinate noise for $\boldsymbol{x}_1$ is projected in this way. The coordinate channels of the other modalities are not independently recentered as the model learns their placement relative to $\boldsymbol{x}_1$.

##### S1.2.2 Denoiser training

For additive Gaussian corruption, Tweedie's formula relates the score $\nabla_{\boldsymbol{X}^{(\sigma)}} \log p_\sigma(\boldsymbol{X}^{(\sigma)})$ of the noised distribution to the conditional mean of the clean sample. We approximate this conditional mean with a denoising network $D_\theta$,

$$\nabla_{\boldsymbol{X}^{(\sigma)}} \log p_\sigma(\boldsymbol{X}^{(\sigma)}) = \frac{\mathbb{E}[\boldsymbol{X}^{(0)} \mid \boldsymbol{X}^{(\sigma)}] - \boldsymbol{X}^{(\sigma)}}{\sigma^2} \approx \frac{D_\theta(\boldsymbol{X}^{(\sigma)}; \sigma) - \boldsymbol{X}^{(\sigma)}}{\sigma^2}. \tag{S4}$$

Thus, estimating the score is equivalent to predicting the clean sample from a noisy observation. The denoiser is trained by minimizing the weighted denoising objective

$$\mathcal{L}(\theta) = \mathbb{E}_{\sigma, \boldsymbol{X}^{(0)}, \boldsymbol{\epsilon}} \left[ \sum_m \lambda_m(\sigma) \left\| D_{\theta,m}(\boldsymbol{X}^{(0)} + \sigma\boldsymbol{\epsilon};\, \sigma) - \boldsymbol{X}_m^{(0)} \right\|_2^2 \right], \qquad \boldsymbol{\epsilon} \sim \mathcal{N}(\boldsymbol{0}, \boldsymbol{I}), \tag{S5}$$

where $m$ indexes the representation variables. The ideal denoiser's optimum is $D^\star(\boldsymbol{X}^{(\sigma)}; \sigma) = \mathbb{E}[\boldsymbol{X}^{(0)} \mid \boldsymbol{X}^{(\sigma)}]$, which ShEPhERD–2 aims to approximate. We therefore use $\hat{\boldsymbol{X}}^{(0)} := D_\theta(\boldsymbol{X}^{(\sigma)}; \sigma)$ as an estimate of the clean molecular structure and interaction profile at any noise level. Following the EDM parameterization [5], we parameterize the denoiser as

$$D_{\theta,m}(\boldsymbol{X}^{(\sigma)}; \sigma) = c_{\text{skip}}(\sigma)\, \boldsymbol{X}^{(\sigma)} + c_{\text{out}}(\sigma)\, F_\theta\Big(c_{\text{in}}(\sigma)\, \boldsymbol{X}^{(\sigma)};\; c_{\text{noise}}(\sigma)\Big), \tag{S6}$$

where $F_\theta$ is a neural network and $c_{\text{skip}}$, $c_{\text{in}}$, $c_{\text{out}}$, and $c_{\text{noise}}$ are $\sigma$-dependent preconditioning coefficients that normalize the input and output scales across noise levels while embedding the noise level itself.

$$\begin{aligned} c_{\text{skip}}(\sigma) &= \frac{\sigma_{\text{data}}^2}{\sigma^2 + \sigma_{\text{data}}^2} & c_{\text{in}}(\sigma) &= \frac{1}{\sqrt{\sigma^2 + \sigma_{\text{data}}^2}}, \\ c_{\text{out}}(\sigma) &= \frac{\sigma\sigma_{\text{data}}}{\sqrt{\sigma^2 + \sigma_{\text{data}}^2}} & c_{\text{noise}}(\sigma) &= \tfrac{1}{4}\log\sigma \\ \lambda(\sigma) &= \frac{1}{c_{\text{out}}(\sigma)^2} \end{aligned} \tag{S7}$$

In Eq. (S6), the scale coefficients act blockwise on the corresponding modality $m$, with indexing suppressed for readability. Equation (S7) gives each block's scalar entries, so $\lambda_m(\sigma) = c_{\text{out},m}(\sigma)^{-2}$. The empirical values of $\sigma_{\text{data},m}$ are reported with the training hyperparameters.

##### S1.2.3 Sampling

At inference time, we traverse the VE noise levels from the Gaussian prior toward the data distribution. The denoising objective specifies the marginal (S2) at each level but does not specify the dependence between states at different levels. For unconditional generation, we use a conditionally independent, non-Markovian coupling,

$$q_{\text{ind}}(\boldsymbol{X}^{(\sigma_1)}, \ldots, \boldsymbol{X}^{(\sigma_N)} \mid \boldsymbol{X}^{(0)}) = \prod_{i=1}^{N} q_{\sigma_i}(\boldsymbol{X}^{(\sigma_i)} \mid \boldsymbol{X}^{(0)}), \tag{S8}$$

which retains the VE marginal in Eq. (S2) at every noise level while using independent noise realizations across levels.

For sampling, we adopt the EDM noise schedule [5],

$$\sigma_i = \left(\sigma_{\min}^{1/\rho} + \frac{i-1}{N-1}\left(\sigma_{\max}^{1/\rho} - \sigma_{\min}^{1/\rho}\right)\right)^{\rho}, \qquad i = 1, \ldots, N, \tag{S9}$$

where $\sigma_0 = 0$ and $\rho$ controls the density of noise levels near $\sigma_{\min}$, so that $\sigma_1 = \sigma_{\min}$ and $\sigma_N = \sigma_{\max}$.

##### S1.2.4 Predict–renoise update

Sampling starts at $\boldsymbol{X}^{(\sigma_N)} \sim \mathcal{N}(\boldsymbol{0}, \sigma_{\max}^2 \boldsymbol{I})$ and iteratively proceeds through decreasing noise levels to $\sigma_{\min}$. At each step, the network first predicts the clean, joint representation and then adds a fresh Gaussian perturbation at the next noise level:

$$\boldsymbol{X}^{(\sigma_{i-1})} = D_\theta(\boldsymbol{X}^{(\sigma_i)}; \sigma_i) + \sigma_{i-1}\boldsymbol{\epsilon}, \qquad \boldsymbol{\epsilon} \sim \mathcal{N}(\boldsymbol{0}, \boldsymbol{I}). \tag{S10}$$

$\boldsymbol{\epsilon}$ is sampled independently at every step, with the noise associated with the atomic coordinates $\boldsymbol{\epsilon}_C$ projected to the COM-less space as in Eq. (S3). Closely related predict–renoise updates have been described as naïve sampling in Cold Diffusion and $\gamma = 1$ sampling in Consistency Trajectory Models [6, 7].

#### S1.3 Conditional generation and inpainting

We partition the representation into a "known" conditioning context, denoted by $\boldsymbol{X}_K$, and the content to be generated, denoted by $\boldsymbol{X}_G$. The context may comprise any subset of representations $\boldsymbol{x}_i$ and/or a subset of their variables $m$. For example, our default interaction profile conditioning strategy sets $\boldsymbol{X}_K = (\boldsymbol{x}_2, \boldsymbol{x}_3, \boldsymbol{x}_4)$ whereas scaffold conditioning specifies only the positions, types, and formal charges ($\boldsymbol{X}_K = (\boldsymbol{C}, \boldsymbol{a}, \boldsymbol{f})$).

##### S1.3.1 Conditional training

We additionally train conditional models in which selected atoms and/or pharmacophores are "fixed" and supplied as clean context $\boldsymbol{X}_F \subseteq \boldsymbol{X}_K$ throughout sampling, enabling scaffold elaboration and pharmacophore prioritization. In this setting, the denoiser is written as $D_\theta(\boldsymbol{X}^{(\sigma)}; \sigma, \boldsymbol{X}_F)$ and is trained to estimate the conditional denoising mean, or equivalently the conditional score, associated with $p(\boldsymbol{X}_G^{(0)} \mid \boldsymbol{X}_F)$.

##### S1.3.2 Inpainting for conditional generation

Before reverse sampling, we independently noise the inpainted context $\boldsymbol{X}_I = \boldsymbol{X}_K \setminus \boldsymbol{X}_F$ at each scheduled noise level. Specifically, we store $\{\bar{\boldsymbol{X}}_I^{(\sigma_i)}\}_{i=0}^N$ where

$$\bar{\boldsymbol{X}}_I^{(\sigma_i)} = \boldsymbol{X}_I^{(0)} + \sigma_i \boldsymbol{\epsilon}_I, \qquad \boldsymbol{\epsilon}_I \sim \mathcal{N}(\boldsymbol{0}, \boldsymbol{I}), \qquad i = 1, \ldots, N. \tag{S11}$$

Thus, each stored state has the appropriate VE marginal distribution, but the noise is independent across noise levels. The content to be generated is initialized from the Gaussian, $\boldsymbol{X}_G^{(\sigma_N)} \sim \mathcal{N}(\mathbf{0}, \sigma_{\max}^2 \boldsymbol{I})$. At each reverse step, the denoiser predicts the complete clean representation,

$$\hat{\boldsymbol{X}}^{(0)} = D_\theta(\boldsymbol{X}^{(\sigma_i)}; \sigma_i). \tag{S12}$$

We retain the content being generated and apply the predict–renoise update to it. The prediction for the inpainted context is discarded and overwritten by the corresponding stored state from the simulated forward trajectory.

$$\boldsymbol{X}_G^{(\sigma_{i-1})} = \hat{\boldsymbol{X}}_G^{(0)} + \sigma_{i-1}\boldsymbol{\epsilon}_G, \qquad \boldsymbol{\epsilon}_G \sim \mathcal{N}(\mathbf{0}, \boldsymbol{I}) \tag{S13}$$

$$\boldsymbol{X}_I^{(\sigma_i)} \leftarrow \bar{\boldsymbol{X}}_I^{(\sigma_i)}. \tag{S14}$$

This repeats until $\sigma_0 = 0$, at which point we reconstruct $\boldsymbol{X}_I^{(0)}$.

##### S1.3.3 Choice of coordinate frame

For conditional generation via inpainting, the molecular structure $\boldsymbol{x}_1$ defines the center of the joint representation, and only its associated sampled noise $\boldsymbol{\epsilon}_C$ is projected to be COM-less. The other modalities retain their placement relative to $\boldsymbol{x}_1$. When the conditioning context fixes a molecular substructure and/or subset of pharmacophoric features, we can instead choose to define the coordinate frame using the COM of the clean context $\boldsymbol{X}_F$ or a slightly perturbed COM of the reference molecule.

##### S1.3.4 Substructure inpainting

For each noise level, we independently noise and store its atomic coordinates, atom types, and bonds, $\{(\boldsymbol{C}_K^{(\sigma_i)}, \boldsymbol{a}_K^{(\sigma_i)}, \boldsymbol{B}_K^{(\sigma_i)})\}_{i=0}^N$. For positions, the sampled noise is centered over these atoms:

$$\tilde{\boldsymbol{\epsilon}}_C = \boldsymbol{\epsilon}_C - \frac{1}{|\mathcal{J}_K|} \sum_{k \in \mathcal{J}_K} \boldsymbol{\epsilon}_{C,k}, \tag{S15}$$

where $\mathcal{J}_K$ is the set of atom indices of the known scaffold. This preserves the COM of the prescribed coordinates without recentering the coordinates themselves. At each reverse step, the prescribed coordinates, atom types, and bonds are overwritten by their stored states at the current noise level (S14), while all unspecified variables continue to be generated. The complete molecular structure is then recentered about the center of mass of its real atoms.

#### S1.4 Composition

Let $\nabla_{\boldsymbol{X}_G^{(\sigma)}} \log p_\sigma(\boldsymbol{X}_G^{(\sigma)} \mid \boldsymbol{X}_K)$ denote the conditional score of the generated structure given a single choice of inpainting $\boldsymbol{X}_K$. With AND composition, we denoise under the joint score defined by $\nabla_{\boldsymbol{X}_G^{(\sigma)}} \log p_\sigma(\boldsymbol{X}_G^{(\sigma)} \mid \boldsymbol{X}_K^1 \ldots \boldsymbol{X}_K^n)$ using the sampling strategy defined in Eq. (S10). Prior compositional diffusion work for image synthesis [8] showed that under assumptions of conditional independence, sampling such a joint score was tractable at inference time. We prove (Suppl. Sec. S3.1.1) that this joint score under (S10) can be expressed as the following AND compositional sampler:

$$\begin{aligned}\boldsymbol{X}_G^{(\sigma_{i-1})} &= D_\theta(\boldsymbol{X}_G^{(\sigma_i)}; \sigma_i) \\ &+ \sum_{c=1}^{n} w_c \Big( D_\theta(\boldsymbol{X}_G^{(\sigma_i)}; \sigma_i, \boldsymbol{X}_K^c) - D_\theta(\boldsymbol{X}_G^{(\sigma_i)}; \sigma_i) \Big) \\ &+ \sigma_{i-1}\boldsymbol{\epsilon}\end{aligned} \tag{S16}$$

$D_\theta(\boldsymbol{X}_G^{(\sigma_i)}; \sigma_i)$ is the unconditional clean-sample prediction and $D_\theta(\boldsymbol{X}_G^{(\sigma_i)}; \sigma_i, \boldsymbol{X}_K^c)$ is the clean-sample prediction under inpainting $\boldsymbol{X}_K^c$, which is weighted by $w_c$.

For NOT composition, we adopt a similar formulation for the joint negation score. We define our favored inpainted interaction profile as $\boldsymbol{X}_{on}$ with any disfavored profile as $\boldsymbol{X}_{off}^c$. Following Liu et al. [8], the negation distribution is formulated as a likelihood ratio, with tilted density $\tilde{p}_\sigma(\boldsymbol{X}) \propto p_\sigma(\boldsymbol{X} \mid \boldsymbol{X}_{on}) \prod_{c=1}^{n} \left( \frac{p_\sigma(\boldsymbol{X}|\boldsymbol{X}_{on})}{p_\sigma(\boldsymbol{X}|\boldsymbol{X}_{off}^c)} \right)^{w_c}$. Sampling this negation joint score $\nabla_{\boldsymbol{X}_G^{(\sigma)}} \log \tilde{p}_\sigma(\boldsymbol{X}_G^{(\sigma)})$ under (S10) is proved in Suppl. Sec. S3.1.2 and defines the NOT compositional sampler:

$$\begin{aligned}\boldsymbol{X}_G^{(\sigma_{i-1})} &= D_\theta(\boldsymbol{X}_G^{(\sigma_i)}; \sigma_i, \boldsymbol{X}_{on}) \\ &+ \sum_{c=1}^{n} w_c \Big( D_\theta(\boldsymbol{X}_G^{(\sigma_i)}; \sigma_i, \boldsymbol{X}_{on}) - D_\theta(\boldsymbol{X}_G^{(\sigma_i)}; \sigma_i, \boldsymbol{X}_{off}^c) \Big) \\ &+ \sigma_{i-1}\boldsymbol{\epsilon}\end{aligned} \tag{S17}$$

#### S1.5 EDM hyperparameters

The hyperparameters used for Suppl. Sec. S1.2-S1.4 are reported in Suppl. Table S1.

#### S1.6 ShEPhERD–2 architecture

These SE(3)-equivariant graph transformer modules individually encode each 3D point cloud into invariant scalar ($\ell = 0$) and equivariant vector ($\ell = 1$) node latent features in the *embedding module*. The *joint module* first processes the joint heterogeneous graph with an EquiformerV3 module that allows the modalities to interact and updates each node embedding. We then sum-pool the $\ell = 1$ embeddings and pass them through an equivariant feedforward network to create an intermediate global $\ell = 1$ representation. This $\ell = 1$ global embedding is combined with an $\ell = 0$ Fourier embedding of the noise level $\sigma$ via an equivariant tensor product. This results in a global invariant and equivariant feature vector which is residually added to the latent node embeddings. Finally, the *denoising module* predicts the fully denoised state of each representation $\hat{\boldsymbol{x}}_i$ where scalar attributes (e.g., atom/pharmacophore types, ESP values) are predicted with multi-layer perceptrons from $\ell = 0$ latent embeddings while positional and vector

**Table S1**: EDM hyperparameters.

| Parameter | Value |
|---|---|
| *Training Parameters* | |
| $P_{\text{mean}}$ | $-1.2$ |
| $P_{\text{std}}$ | 1.2 |
| $\sigma_{\text{data}}$ (positions) | 3.0 |
| $\sigma_{\text{data}}$ (ESP) | 0.46 |
| $\sigma_{\text{data}}$ (atom one-hot) | 0.3 |
| $\sigma_{\text{data}}$ (pharm one-hot) | 0.27 |
| $\sigma_{\text{data}}$ (bonds) | 0.43 |
| *Inference Parameters* | |
| $\sigma_{\text{max}}$ | 3.0 |
| $\sigma_{\text{min}}$ | 0.001 |
| $\rho$ | 7.0 |
| $N$ | 400 |
| *Composition Parameters* | |
| $w_c$ | 1/n |

attributes (e.g., atom positions, pharmacophore vectors) are predicted with equivariant feedforward networks [9, 10] from $\ell = 1$ latent embeddings. Technical details of these modules are explained in depth in Adams et al. [11] and key parameters are described in Suppl. Table S2.

#### S1.7 ShEPhERD–2 training

We augment each training sample by adding in $n_{\text{da}} \sim \mathcal{U}(0, \min(10, n_1))$ dummy atoms and $n_{\text{dp}} \sim \mathcal{U}(0, \min(5, n_4))$ dummy pharmacophores. The positions of randomly selected atoms (without replacement) are used as the coordinates for the dummy atoms after adding isotropic Gaussian noise $\boldsymbol{\xi} \sim \mathcal{N}(\mathbf{0}, \sigma_{\text{da}}^2 \boldsymbol{I})$ where $\boldsymbol{\xi} \in \mathbb{R}^{n_{\text{da}} \times 3}$. We apply the same approach for dummy pharmacophore positions by using the noised coordinates of randomly selected pharmacophore features. The dummy pharmacophore vector is either directionless $\boldsymbol{v}_{\text{dp}} = \mathbf{0}$ with a probability $p_{\text{dp}}$, or is a unit vector $\boldsymbol{v}_{\text{dp}} = \frac{\boldsymbol{\nu}}{\|\boldsymbol{\nu}\|}$ where $\boldsymbol{\nu} \sim \mathcal{N}(\mathbf{0}, \boldsymbol{I})$ with a probability $1 - p_{\text{dp}}$.

For each training example, we sample a noise magnitude $\sigma$ according to the EDM parameterized log-normal distribution $\ln(\sigma) \sim \mathcal{N}(P_{\text{mean}}, P_{\text{std}}^2)$ [5]. Independent Gaussian noise $\boldsymbol{\epsilon}_m \sim \mathcal{N}(\mathbf{0}, \sigma^2 \boldsymbol{I})$ is then added to each representation according to the variance-exploding (VE) noising process in Eq. (S1). The preconditioned denoiser in Eq. (S6) is trained to predict the clean sample from the resulting noisy sample using the weighted L2 loss in Eq. (S5). Training hyperparameters are in Suppl. Table S3.

We additionally train the model to condition generation on fixed atoms, substructures, and pharmacophore features. With probability $p_{\text{fa}}$, a subset of the molecular structure $\boldsymbol{x}_{1,F}$ is fixed. This subset consists of either (1) up to $n_{\text{fa}} \leq n_1$ randomly selected atoms, with hydrogen atoms selected at a reduced probability, or (2) up to four randomly selected connected substructures with up to 9 atoms each. The corresponding atomic coordinates, types, and formal charges remain unnoised during forward noising. Similarly, with probability $p_{\text{fp}}$, a subset of pharmacophore features $\boldsymbol{x}_{4,F}$ is fixed, retaining their types, positions, and directions throughout the diffusion trajectory. In this fixed *conditional* setting, the coordinate frame is either placed at a

**Table S2**: Architecture hyperparameters.

| ***Denoising Network Hyperparameters*** | |
|---|---|
| **Parameter** | **Value** |
| ***Default EquiformerV3 Parameters*** | |
| num_node_channels | 64 |
| lmax_list | [1] |
| mmax_list | [1] |
| ffn_hidden_channels | 32 |
| grid_resolution | 4 |
| num_sphere_samples | 128 |
| edge_channels | 128 |
| attn_activation | sep-merge_gates2_swiglu |
| use_add_merge | False |
| use_grid_mlp | True |
| ***Joint Module Parameters*** | |
| num_EquiformerV3_layers | 2 |
| attention_channels | 24 |
| num_attention_heads | 2 |
| radius_graph_cutoff | 5.0 |
| RBF_cutoff | 5.0 |
| $\boldsymbol{x}_1$ ***Embedding Module Parameters*** | |
| num_EquiformerV3_layers | 4 |
| attention_channels | 32 |
| num_attention_heads | 4 |
| ffn_hidden_channels | 64 |
| radius_graph_cutoff | $\infty$ (fully connected) |
| RBF_cutoff | 5.0 |
| $\boldsymbol{x}_3$, $\boldsymbol{x}_4$ ***Embedding Module Parameters*** | |
| num_EquiformerV3_layers | 2 |
| attention_channels | 24 |
| num_attention_heads | 2 |
| ffn_hidden_channels | 32 |
| radius_graph_cutoff | 5.0 |
| RBF_cutoff | 5.0 |
| x3_scalar_RBF_expansion_min | -10.0 |
| x3_scalar_RBF_expansion_max | 10.0 |
| $\boldsymbol{x}_1$ ***Denoising Module Parameters*** | |
| MLP_hidden_dim | 64 |
| num_MLP_hidden_layers | 2 |
| e3nn_ffn_hidden_channels | 32 |
| egnn_normalize_vectors | True |
| egnn_distance_expansion_dim | 32 |
| $\boldsymbol{x}_3$, $\boldsymbol{x}_4$ ***Denoising Module Parameters*** | |
| MLP_hidden_dim | 64 |
| num_MLP_hidden_layers | 2 |
| e3nn_ffn_hidden_channels | 32 |

**Table S3**: Training hyperparameters.

| *Training hyperparameters* | |
|---|---|
| **Parameter** | **Value** |
| $n_2$ | 75 |
| surface probe radius | 0.6 |
| $\boldsymbol{a}$ and $\boldsymbol{f}$ scaling factors | 0.25 |
| $\sigma_{\text{da}}$ (dummy atom noise std) | 0.2 |
| $p_{\text{dp}}$ (directional dummy pharm.) | 0.500 |
| $p_{\text{fa}}$ (fix atoms only) | 0.167 |
| $p_{\text{fp}}$ (fix pharm. only) | 0.167 |
| $p_{\text{fa,fp}}$ (fix both) | 0.167 |
| ***Optimization parameters*** | |
| Effective batch size | 96 |
| Learning rate | 0.001 |
| Minimum learning rate | 0.0003 |
| Warmup steps | 10000 |
| Learning rate decay steps | 100000 |
| Gradient clipping value | 5.0 |

reference's slightly noised COM or at the COM of the clean context $\boldsymbol{X}_F$. In the case of the latter, we do not project $\boldsymbol{\epsilon}_C$ to be COM-less. Dummy atoms or pharmacophore features are prevented from being placed on the fixed context, and we mask out the fixed features from contributing to the loss.

We trained ShEPhERD–2 for 300k optimization steps using the Adam optimizer with an effective batch size of 96 on four NVIDIA H200 which took approximately 2 days. We use FP32 precision during training.

#### S1.8 3D similarity scoring

Detailed explanations of the scoring functions and their parameterization are found in Adams et al. [11]; we summarize them below. Given a point cloud-based molecular representation, we place isotropic Gaussians at the coordinates of the sample point cloud $\boldsymbol{Q}_A \in \mathbb{R}^{n_i \times 3}$ for $\boldsymbol{x}_i$ where $i$ is a specified interaction modality with $n_i$ points. The first-order Gaussian overlap between $\boldsymbol{Q}_A$ and the reference $\boldsymbol{Q}_B$ is

$$O_{AB} = \sum_{a \in A} \sum_{b \in B} w_{ab} \frac{\pi^{3/2}}{(2\alpha)^{3/2}} \exp\left(-\frac{\alpha}{2} \left\|\mathbf{r}_a - \mathbf{r}_b\right\|^2\right), \tag{S18}$$

where $\alpha$ is the Gaussian width associated with the identity of the point representation and $w_{ab}$ is a weighting function that accounts for differences in ESP or vector directionality of matching pharmacophore features [12–16]. The Tanimoto similarity of the overlaps is

$$\text{sim}(\boldsymbol{Q}_A, \boldsymbol{Q}_B) = \frac{O_{AB}}{O_{AA} + O_{BB} - O_{AB}}. \tag{S19}$$

We report the optimal 3D similarity, which is found by SE(3) rigid transformations of $\boldsymbol{Q}_A$ to maximize the similarity to $\boldsymbol{Q}_B$:

$$\text{sim}^*(\boldsymbol{Q}_A, \boldsymbol{Q}_B) = \max_{\boldsymbol{R},\boldsymbol{t}} S(\boldsymbol{R}\boldsymbol{Q}_A + \boldsymbol{t}, \boldsymbol{Q}_B), \tag{S20}$$

where $\boldsymbol{R} \in SO(3)$ and $\boldsymbol{t} \in T(3)$. We typically conduct *local* alignment which optimizes this function starting only from the sampled (post-relaxation) pose for up to 200 steps with the Adam optimizer. If specified, we also conduct *global* optimization, which simultaneously aligns many randomly seeded poses and reports the maximum score and associated pose.

To compute the 3D similarity of a *subset* of an interaction profile (e.g., high priority or low priority pharmacophore), we utilize Tversky similarity $\text{sim}_{\text{tversky}}(\boldsymbol{Q}_A, \boldsymbol{Q}_B) = \frac{O_{AB}}{O_{BB}}$ computed after alignment to the *full* profile (S20).

**Shape similarity**. The shape similarity between two surfaces $\boldsymbol{S}_A$ and $\boldsymbol{S}_B$ is $\text{sim}^*_{\text{surf}}(\boldsymbol{S}_A, \boldsymbol{S}_B)$ where $\alpha = 1.258$ and $w_{A,B} = 1$.

**Electrostatic similarity**. The similarity between two electrostatic surfaces $\boldsymbol{x}_{3,A}$ and $\boldsymbol{x}_{3,B}$ is $\text{sim}^*_{\text{ESP}}(\boldsymbol{x}_{3,A}, \boldsymbol{x}_{3,B})$ where $\alpha = 1.258$, $w_{A,B} = \frac{\|v_{A,a} - v_{B,b}\|^2}{\lambda}$, and $\lambda = \frac{0.3}{(4\pi\epsilon_0)^2}$. Here, $\epsilon_0$ is the permittivity of vacuum.

**Pharmacophore similarity**. The similarity between two pharmacophores $\boldsymbol{x}_{4,A}$ and $\boldsymbol{x}_{4,B}$ is

$$\text{sim}^*_{\text{pharm.}}(\boldsymbol{x}_{4,A}, \boldsymbol{x}_{4,B}) = \frac{\sum_{m \in \mathcal{M}} O_{A,B;m}}{\sum_{m \in \mathcal{M}} O_{A,A;m} + O_{B,B;m} - O_{A,B;m}},$$

where $\mathcal{M}$ is the set of all pharmacophore types, the Gaussian width $\alpha_m$ is dependent on the pharmacophore type as defined by Taminau et al. [17], and the weight is implemented following Wahl [16]

$$w_{a,b;m} = \begin{cases} 1 & \text{if } m \text{ is directionless,} \\ \frac{\boldsymbol{V}^\top_{A,a;m} \boldsymbol{V}_{B,b;m} + 2}{3} & \text{if } m \text{ is directional.} \end{cases}$$

We assume that the $\pi$ effects of aromatic groups are symmetric across the plane and take the absolute value of $\boldsymbol{V}^\top_{A,a;m} \boldsymbol{V}_{B,b;m}$.

#### S1.9 Default sampling settings

We use the EDM noising schedule for sampling (S9), starting from $\sigma_{\max} = 3.0$ and progressively decreasing to $\sigma_{\min} = 0.001$ with $\rho = 7$ and $N = 400$ (Suppl. Table S1). We terminate sampling after 360 sampling steps because predictions converge with no observable loss in quality, reducing the computational cost by 10%. We further accelerate sampling using TensorFloat-32 (TF32) Tensor Cores via `torch.set_float32_matmul_precision('high')`, with no measured loss in quality.

##### S1.10 MolGenBench hit-to-lead benchmark

The reported metrics and their definitions are explained in Suppl. Table S17. The protocol used for each baseline model, an explanation of the different validity metrics, and uncertainty quantification with bootstrapping are detailed below.

###### S1.10.1 Baseline model protocols

**FLOWR.ROOT** [18]. FLOWR.ROOT (commit hash: 9ff49e8) applies interaction-conditioned generation by inpainting atoms involved in interactions. In the ligand-based setting, we use the `--scaffold_hopping` flag, which inpaints atoms outside the Murcko scaffold (e.g., peripheral substituents). In the structure-based setting, we use the `--interaction_conditional` flag to inpaint atoms involved in interactions identified by ProLIF while explicitly conditioning on the atomic geometry of the protein pocket.

**PhoreGen** [19]. PhoreGen (commit hash: 6f3f7f0) applies AncPhore [20] identified pharmacophores and exclusion spheres as fixed conditioning during diffusion. PhoreGen does not natively support the ligand-based setting, so we randomly subselect ligand pharmacophores and exclusion sphere positions consistent with PhoreGen's synthetic-ligand-only pretraining protocol. In the structure-based setting, we use the `train_dock-cpx-phore` model in `sample_all` mode to condition on the identified protein-ligand pharmacophore model and exclusion spheres based on protein shape.

**OMTRA** [21]. OMTRA (commit hash: 745127d) generates molecules conditioned on directionless pharmacophores extracted from a reference ligand. For the ligand-based setting, we use the `denovo_ligand_from_pharmacophore_condensed` mode, which utilizes the `omtra-v0-pharm-denovo` model. For pocket-conditioned design, the `fixed_protein_pharmacophore_ligand_denovo_condensed` mode conditions generation on the same ligand pharmacophore and explicit pocket atom geometry using the `omtra-v0-protpharm-denovo` model.

**CondSemla** [22]. CondSemla (commit hash: 22d5159) generates interaction-conditioned molecules by keeping hydrogen-bond acceptor, donor, and aromatic atoms unnoised throughout inference. For this ligand-based setting, we use the default `replacement_guidance`, which utilizes the `geom-drugs-200epochs` model from SemlaFlow [23].

**DiffDec** [24] and **DiffSBDD** [25]. We utilize the samples provided by MolGenBench [26] for these models to compute the MolGenBench metrics. Specifically, we report the version of DiffSBDD trained on CrossDocked because it reported improved interaction scores, Vina affinity, and diversity metrics despite slightly lower validity. These models condition directly on a specified fragment derived from the reference compound. DiffDec is trained to condition on a fixed substructure whereas DiffSBDD inpaints the structure.

###### S1.10.2 Validity metrics

We distinguish molecular validity as reported elsewhere from the MolGenBench validity metrics. "Conformer Validity" is the PoseBusters (PB) intramolecular validity, which requires a non-relaxed sample to yield an RDKit-parseable molecular graph and satisfy heuristic checks on molecular geometry (e.g., bond lengths and angles) and global strain as approximated by the "PB Energy Ratio". "PB valid" includes all "Conf. Validity" criteria and additionally requires that the molecule does not clash with the pocket [27].

##### S1.10.3 Uncertainty estimation and statistical testing

Confidence intervals are estimated using a cluster bootstrap over UniProt protein targets, with all molecules and series from each sampled target kept together. Metrics are recomputed for each bootstrap sample, and the 2.5th and 97.5th percentiles define the 95% confidence interval. Point estimates are unchanged. For model comparisons, we use two-sided paired tests on 10,000 cluster-bootstrap resamples of the same target sets, with pairwise differences evaluated using step-down max-$t$ multiplicity correction across models within each table row.

#### S1.11 Interaction-aware genetic algorithm

Let $\mathcal{P}_0 = \{\mathcal{I}^{(k)}\}_{k=1}^{K}$ denote the intial population of individuals $\mathcal{I} = (\boldsymbol{X}, s)$ where $\boldsymbol{X} = (\boldsymbol{x}_1, \boldsymbol{x}_2, \boldsymbol{x}_3, \boldsymbol{x}_4)$ is the complete profile with $\boldsymbol{X}_K = (\boldsymbol{x}_2, \boldsymbol{x}_3, \boldsymbol{x}_4)$ referring to the interaction channels and $\boldsymbol{x}_1$ is the 3D molecular structure. $s$ refers to a scalar or n-dimensional tuple representing an individual's fitness. With some abuse of notation, the interaction-aware genetic algorithm seeks the individual $\mathcal{I}^\star$ of highest fitness $s$ reachable from the heritable set of interactions of $\mathcal{P}_0$

$$\mathcal{I}^\star = \underset{\substack{\boldsymbol{x}_1 \in \mathcal{D}_\theta(\boldsymbol{X}_K) \\ \boldsymbol{X}_{K_0} \in \mathcal{P}_0}}{\arg\max}\; s(\mathcal{I}_{\boldsymbol{x}_1}). \tag{S21}$$

where $\mathcal{D}_\theta(\boldsymbol{X}_K)$ refers to the general chemical space accessible by ShEPhERD–2 given an interaction profile and $\boldsymbol{X}_{K_0}$ is the set of seed interaction profiles. The genotype space for heuristic mutation and crossover operators is therefore interaction profiles, while the phenotype space is the molecular structure.

##### S1.11.1 Interaction mutation and crossover

With probability $1 - p_\times$, an offspring is produced by mutation where a single parent interaction profile $\boldsymbol{X}_K$ supplies the inpainting context (Sec. S1.3). Unless stated otherwise, all interaction channels are inpainted. The atom and pharmacophore counts of the offspring are sampled uniformly from ranges $N_{x_1} \sim \max\big(1, \mathcal{U}\{N^{\text{ref}}_{\text{atoms}}+l_1,\, N^{\text{ref}}_{\text{atoms}}+h_1\}\big)$, $N_{x_4} \sim \max\big(1, \mathcal{U}\{N^{\text{ref}}_{\text{pharm}},\, N^{\text{ref}}_{\text{pharm}}+h_4\}\big)$, with lower and upper offsets $l_1, h_1, h_4$.

With probability $p_\times$, an offspring is produced by crossover of two parents where profiles $a, b$ are selected and the AND-compositional sampler of Eq. (S16) is applied over inpainting context $\boldsymbol{X}^a_K$ and $\boldsymbol{X}^b_K$. Each profile pair is re-centered according to the midpoint of their two centers of mass and composed under weights $w_a, w_b$. $N^{\text{ref}}_{\text{atoms}}$ and $N^{\text{ref}}_{\text{pharm}}$ are defined as $\max(N_a, N_b)$ and otherwise follow the same count formulation as mutation. Optionally, fragment merging exploration can be enabled by $N_a + N_b$ if both paired structures' heavy-atom counts are below the fragment threshold $\tau$, or by $N_a + \alpha \min(N_a, N_b)$ if only one of the pair's heavy-atom counts is below $\tau$.

All samples are post-relaxed with GFN2-xTB, and RDKit-valid molecules are then subjected to user-defined constraints, including drug-likeness and assay promiscuity. Samples passing all these criteria are added to the generation's population $\mathcal{P}$. After scoring individuals $\mathcal{I}$, some oracles output predicted conformers associated with the score $s$ (e.g., a predicted bound pose); in such cases, we defer to the oracle-chosen conformer and re-extract the interaction profile for subsequent generations.

##### S1.11.2 Selection

We determine which individuals of $\mathcal{P}$ act as parents for interaction mutation and crossover based on fitness $s$. Because $s(\mathcal{I}_{\boldsymbol{x}_1})$ may be scalar or an $n$-tuple, we define both single-objective and Pareto ranking.

For Pareto ranking, we adopt the NSGA-II algorithm [28]. Briefly, an individual $\mathcal{I}_i$ *dominates* $\mathcal{I}_j$, $\mathcal{I}_i \succ \mathcal{I}_j$, if it is at least as good on every objective and strictly better on at least one. Repeated non-dominated sorting of $\mathcal{P}$ under $\succ$ partitions the population into fronts $\mathcal{F}_0, \mathcal{F}_1, \ldots$, with $\mathcal{F}_0 = \{\mathcal{I} \in \mathcal{P} : \nexists \mathcal{I}' \in \mathcal{P}, \mathcal{I}' \succ \mathcal{I}\}$ and each subsequent $\mathcal{F}_k$ is defined identically after removing $\bigcup_{j<k} \mathcal{F}_j$; every individual is assigned the Pareto rank $\rho(\mathcal{I}) = k$ of the front $\mathcal{F}_k$ it belongs to, and tiebreaks are based on NSGA-II crowding distance.

Parents are drawn by tournament selection based on this Pareto rank $\rho(\mathcal{I})$ or scalar fitness $s$: a subset $\mathcal{T} \subset \mathcal{Q}$ of size $T$ is sampled uniformly without replacement from a parent pool $\mathcal{Q}$, and the best-ranking individual $\mathcal{I} \in \mathcal{T}$ is selected. $\mathcal{Q}$ is defined as either the current population $\mathcal{P}$ at generation $g$ or a persistent top-$k$ archive maintained across generations. If the latter is enabled and $s$ is multiobjective, we recalculate Pareto ranks with respect to this top-$k$ pool.

##### S1.11.3 Bioisosteric fragment merging genetic algorithm parameters

We outline the respective parameters used during the exploration and exploitation phases of bioisosteric fragment merging for EV-D68 3C protease in Suppl. Table S4.

#### S1.12 Sub-selection of negation interaction profile

We trilinearly interpolate the energy grid Vina constructs for each atom type to score heavy atoms at every position. This returns the Vina first-order energy decomposition of each heavy atom, whose summation is the total Vina docking score. For negation considered in this work, the reference molecule has the same number of heavy atoms in its on- and off-target pose, thus allowing us to calculate the energy difference atom-wise. Absolute differences above 0.1 in favor of the off-target structure are assumed differentially favorable for the off-target and define a negation atom set. All pharmacophores $\boldsymbol{x}_4$ associated with negation atoms are included in the negation profile. $\boldsymbol{x}_2$ shape and $\boldsymbol{x}_3$ electrostatics are similarly constructed from all atoms that are associated with these negated pharmacophores, including explicit hydrogens.

**Table S4**: Genetic algorithm hyperparameters

| ***GA Hyperparameters*** | | |
|---|---|---|
| **Parameter** | **Exploration** | **Exploitation** |
| ***Population Parameters*** | | |
| population_size | 50 | 50 |
| num_generations | 20 | 20 |
| max_iterations_fraction | 4.0 | 10.0 |
| top_k | 50 | 50 |
| ***Selection Parameters*** | | |
| selection | tournament | tournament |
| tournament_size | 8 | 8 |
| ***Crossover Parameters*** | | |
| crossover_w | 0.5, 0.5 | 0.5, 0.5 |
| crossover_prob | 0.8 | 0.3 |
| crossover_batch_size | 4 | 4 |
| ***Mutation Parameters*** | | |
| mutation_mode | inpainting | inpainting |
| condition_mode | all | all |
| N_x1_range | 0, 6 | 0, 6 |
| N_x4_range | 0, 2 | 0, 2 |
| mutate_batch_size | 4 | 4 |
| ***Fragment-Merge Parameters*** | | |
| fragment_atom_threshold | 25 | N/A |
| fragment_inclusion_prob | 0.3 | N/A |
| fragment_merge_alpha | 0.4 | N/A |
| ***Validity Filter Parameters*** | | |
| validity | composite | composite |
| mw_limit | 500 | 500 |
| logp_limit | 5.0 | 5.0 |
| hbd_limit | 5 | 5 |
| hba_limit | 10 | 10 |
| max_violations | 1 | 1 |
| rotatable_bonds_limit | 10 | 10 |
| sa_score_limit | 4.5 | 4.5 |
| promiscuity_filters | pains, brenk | pains, brenk |

### S2 Extended Data

#### S2.1 ShEPhERD–2 improvements

Replacing the EquiformerV2 modules with EquiformerV3 and adopting the variance-exploding formulation in ShEPhERD–2 yields a 1.8× speedup relative to ShEPhERD. Additional code optimizations further improve runtime and memory efficiency, as reported in Suppl. Table S6. Note that these were measured at a batch size of 20, but the improvements in ShEPhERD–2 enable much larger batch sizes (e.g., 192 for $n_1 = 50$ on an H200), which can generate at nearly twice the throughput.

**Table S5**: Ablation of ShEPhERD–2 design choices on the MOSES test set ($N^{ref}_{atom/pharm}$). Each row adds a component to the row above where the additions of dummy atoms/pharmacophores, EDM-style variance exploding diffusion [5], and EquiformerV3 (EqV3) [29] lead to ShEPhERD–2. Validity is the post-xTB-optimization validity rate (%) with a 95% confidence interval. Strain energy (kcal/mol) is the median over all valid molecules. Diversity is the median post-optimization graph similarity to reference (lower is better). Shape, ESP, and pharmacophore similarities are the median best score among graph-diverse samples (graph similarity $< 0.3$) per reference molecule. The best value in each column is shown in bold while the second best is underlined.

| Configuration | Validity (%) [95% CI] | Strain (kcal/mol) | Diversity | Shape | ESP | Pharm. |
|---|---|---|---|---|---|---|
| ShEPhERD | 61.3 [56.5, 66.1] | 0.052 | **0.165** | 0.843 | 0.836 | 0.619 |
| + dummy atoms/pharm. | 84.7 [81.9, 87.4] | 0.035 | 0.195 | 0.847 | 0.837 | 0.670 |
| + EDM | **94.0** [92.5, 95.4] | <u>0.026</u> | <u>0.170</u> | <u>0.863</u> | <u>0.858</u> | <u>0.718</u> |
| + EqV3 (ShEPhERD–2) | <u>93.7</u> [92.4, 95.0] | **0.015** | 0.189 | **0.864** | **0.860** | **0.721** |

**Table S6**: Comparison of ShEPhERD vs. ShEPhERD–2 efficiency. Forward pass, backpropagation, and memory are measured on a single NVIDIA H100 80GB HBM3 with a batch size of 20. Forward-pass speed and forward memory are measured with a `no_grad` inference forward. Training memory is the peak over a full training step (forward + backward + optimization). Per-step values are mean ± std over 100 steps. Note that because the memory footprint is lower with ShEPhERD–2, we can also increase the batch size whereas ShEPhERD is limited by memory. We also report the time required to run the MOSES evaluation at ($N^{ref}_{atom/pharm}$) on NVIDIA H200 141GB HBM3e NVL. ShEPhERD–2 requiring 10% fewer steps during a denoising trajectory, PyTorch compilation caching, and general code refinements explain the additional speedup when comparing per-step to batch.

| | **ShEPhERD** | **ShEPhERD–2** | **Improvement** |
|---|---|---|---|
| *Per step* (H100) | | | |
| Forward pass (ms) | $279.1 \pm 4.1$ | $56.6 \pm 2.0$ | 4.93× |
| Backprop (ms) | $221.6 \pm 4.8$ | $68.3 \pm 1.8$ | 3.24× |
| Forward memory (GB) | $43.75 \pm 0.54$ | $0.99 \pm 0.02$ | 44.2× |
| Training memory (GB) | $58.86 \pm 1.20$ | $9.65 \pm 0.27$ | 6.0× |
| *MOSES test set* (H200) | | | |
| Full test set (hr) | 3.05 | 0.48 | 6.35× |
| Per batch (s) | 109.8 | 17.3 | 6.35× |

#### S2.2 MOSES test set

##### S2.2.1 Comparison between ShEPhERD and ShEPhERD–2

**Table S7**: Fraction valid after xTB relaxation reported as the mean and 95% CI.

| Model | Atom / pharmacophore count relative to reference | | |
|---|---|---|---|
| | $N^{\text{ref}}_{\text{atoms}}$ $N^{\text{ref}}_{\text{pharm}}$ | $N^{\text{ref}}_{\text{atoms}}+4$ $N^{\text{ref}}_{\text{pharm}}+2$ | $N^{\text{ref}}_{\text{atoms}}+8$ $N^{\text{ref}}_{\text{pharm}}+4$ |
| ShEPhERD | 0.613 [0.565, 0.661] | 0.497 [0.460, 0.534] | 0.413 [0.378, 0.448] |
| ShEPhERD–2 | **0.937** [0.924, 0.950] | **0.942** [0.928, 0.956] | **0.937** [0.924, 0.949] |

**Table S8**: Median benchmark results for ShEPhERD vs. ShEPhERD–2. Statistical comparisons are two-sided Mann–Whitney U tests, Holm-corrected within each block. For strain energy, lower is better (positive Cliff's $\delta$ favors ShEPhERD–2). For 3D similarity and QED, higher is better (negative Cliff's $\delta$ favors ShEPhERD–2). The better median in each row is shown in bold.

| Quantity | Setting | Median | | $p$ | Cliff's $\delta$ |
|---|---|---|---|---|---|
| | | ShEPhERD | ShEPhERD–2 | | |
| *Strain energy (kcal/mol)* | | | | | |
| Strain | $N^{\text{ref}}_{\text{atoms}}$, $N^{\text{ref}}_{\text{pharm}}$ | 0.052 | **0.015** | $3.2\times10^{-249}$ | +0.660 |
| Strain | $N^{\text{ref}}_{\text{atoms}}+4$, $N^{\text{ref}}_{\text{pharm}}+2$ | 0.065 | **0.016** | $<10^{-300}$ | +0.763 |
| Strain | $N^{\text{ref}}_{\text{atoms}}+8$, $N^{\text{ref}}_{\text{pharm}}+4$ | 0.090 | **0.018** | $<10^{-300}$ | +0.801 |
| *QED* | | | | | |
| QED | $N^{\text{ref}}_{\text{atoms}}$, $N^{\text{ref}}_{\text{pharm}}$ | 0.773 | **0.798** | $1.1\times10^{-9}$ | −0.129 |
| QED | $N^{\text{ref}}_{\text{atoms}}+4$, $N^{\text{ref}}_{\text{pharm}}+2$ | 0.766 | **0.801** | $1.1\times10^{-16}$ | −0.190 |
| QED | $N^{\text{ref}}_{\text{atoms}}+8$, $N^{\text{ref}}_{\text{pharm}}+4$ | 0.736 | **0.801** | $1.7\times10^{-46}$ | −0.347 |
| *3D similarity — (graph similarity < 0.3)* | | | | | |
| Shape | all | 0.740 | **0.769** | $7.7\times10^{-11}$ | −0.151 |
| ESP | all | 0.708 | **0.751** | $1.4\times10^{-16}$ | −0.194 |
| Pharm. | all | 0.475 | **0.524** | $1.4\times10^{-16}$ | −0.195 |
| Shape | top-1 | 0.843 | **0.864** | $2.8\times10^{-4}$ | −0.299 |
| ESP | top-1 | 0.836 | **0.860** | $2.4\times10^{-5}$ | −0.359 |
| Pharm. | top-1 | 0.619 | **0.721** | $1.7\times10^{-8}$ | −0.478 |

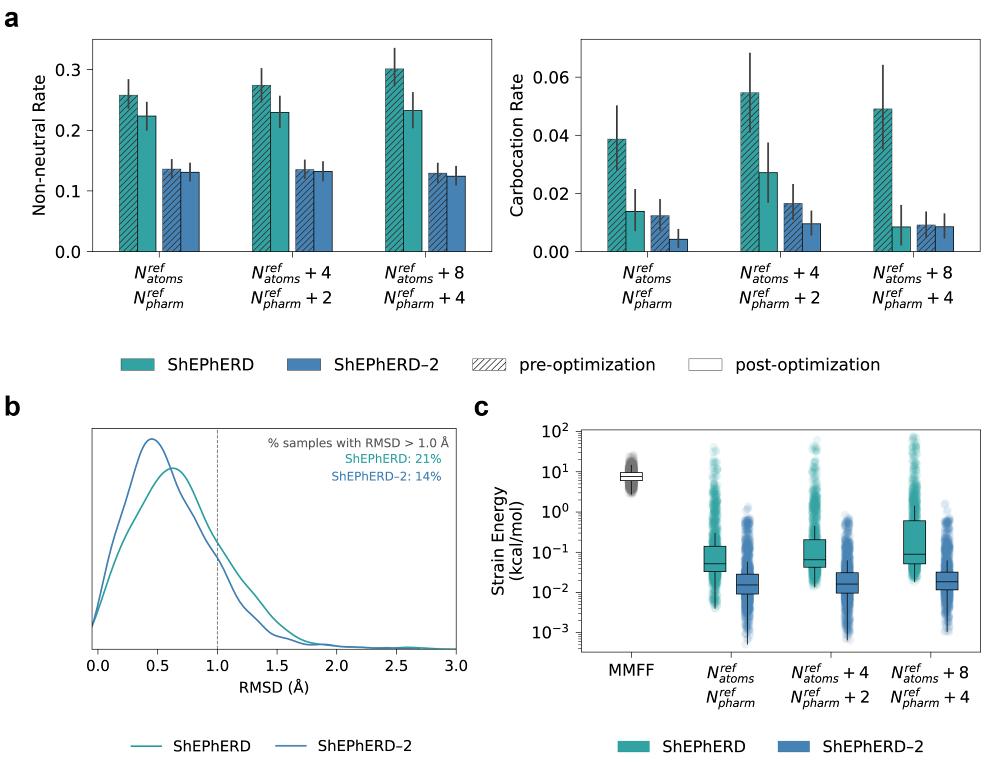

**Fig. S1**: **a**, The rate of sampling non-neutral species or carbocations pre-/post-xTB optimization. **b**, Heavy-atom root-mean-square deviation (RMSD) after geometry relaxation with GFN2-xTB. More than 20% of samples generated by ShEPhERD exhibit RMSD values greater than 1.0 Å. In contrast, fewer than 15% of ShEPhERD–2-generated samples exceed 1.0 Å RMSD. **c**, Strain energy (kcal/mol) distributions following local GFN2-xTB relaxation for conformers generated by ShEPhERD and ShEPhERD–2. The reference distribution was computed from 2,000 randomly selected MOSES training-set molecules that were initially optimized with MMFF and subsequently re-optimized with GFN2-xTB.

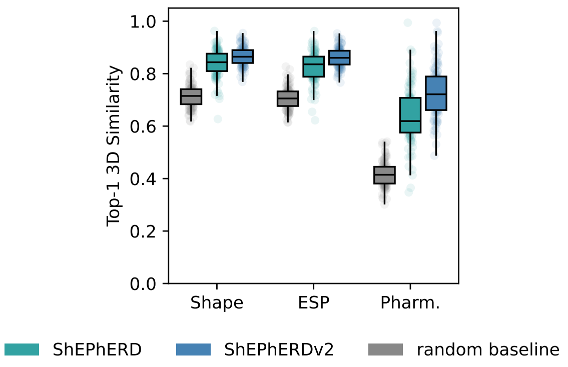

**Fig. S2**: Distributions of 3D similarities of the best-scoring sample per MOSES test set reference with graph similarity < 0.3. ShEPhERD–2 significantly enriches these distributions relative to ShEPhERD (Suppl. Table S8).

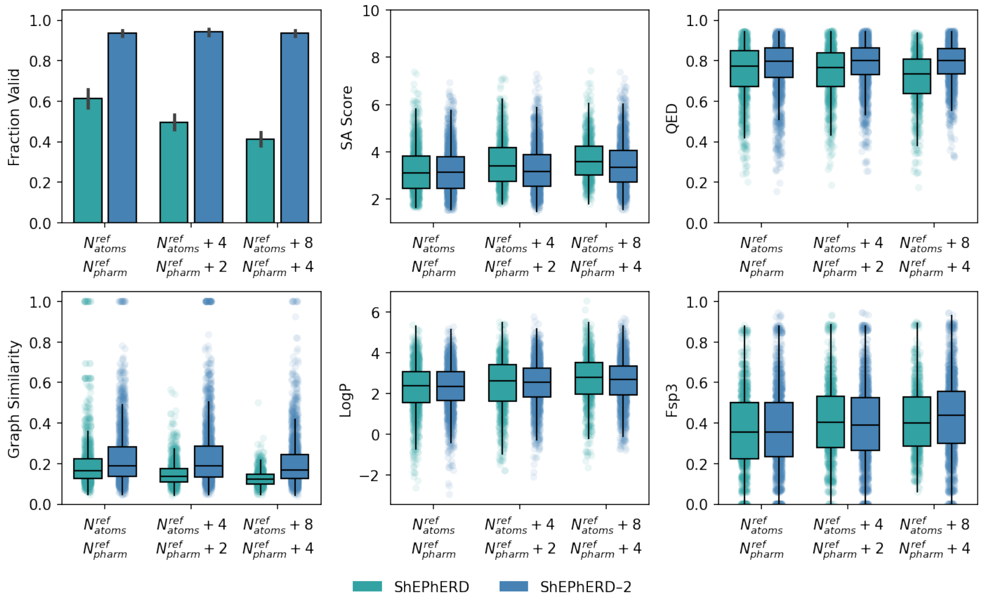

**Fig. S3**: Sample quality of generated samples (post-relaxation) at increasing number of atoms and pharmacophores relative to the reference.

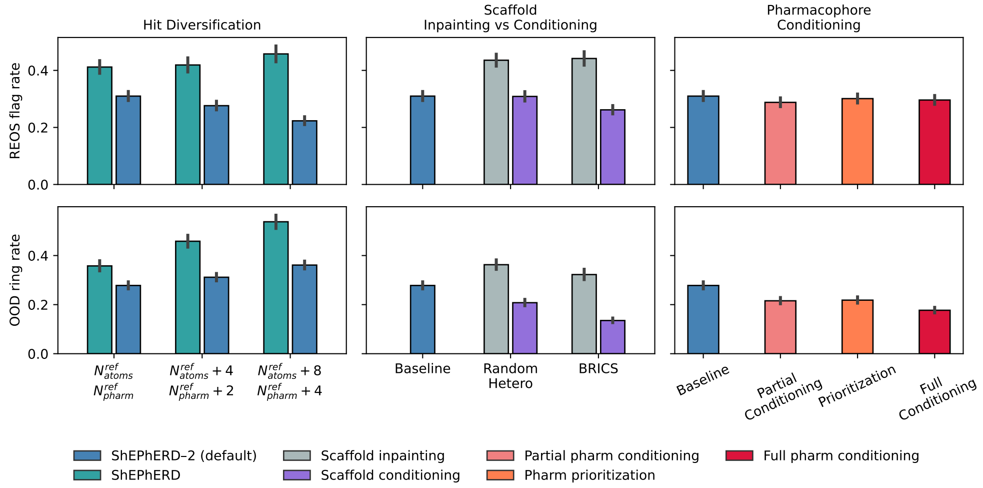

**Fig. S4**: **(Top)** The rate of sampling molecules with at least one REOS (rapid elimination of swill) flag after relaxation [30]. **(Bottom)** The rate of sampling molecules with at least one out-of-distribution (OOD) ring system that does not have precedent in ChEMBL [31, 32].

##### S2.2.2 Pharmacophore prioritization

**Table S9**: Pairwise pharmacophore-recovery comparisons across four conditioning strategies on the MOSES test set (graph similarity $< 0.3$). Statistical comparisons are two-sided Mann–Whitney U tests, Holm-corrected within each feature set. Higher similarity is better; a positive Cliff's $\delta$ favors the first-named strategy (A). The larger median in each row is shown in bold.

| Comparison (A vs. B) | Median | | $p$ | Cliff's $\delta$ |
|---|---|---|---|---|
| | A | B | | |
| *Full pharmacophore set* | | | | |
| standard vs. HP only | **0.524** | 0.506 | 0.002 | +0.066 |
| standard vs. prioritization | 0.524 | **0.574** | $1.2 \times 10^{-17}$ | −0.191 |
| standard vs. all fixed | 0.524 | **0.627** | $2.3 \times 10^{-44}$ | −0.339 |
| HP only vs. prioritization | 0.506 | **0.574** | $4.5 \times 10^{-32}$ | −0.255 |
| HP only vs. all fixed | 0.506 | **0.627** | $6.3 \times 10^{-62}$ | −0.393 |
| prioritization vs. all fixed | 0.574 | **0.627** | $1.1 \times 10^{-10}$ | −0.161 |
| *High-priority pharmacophores* | | | | |
| standard vs. HP only | 0.701 | **0.799** | $1.7 \times 10^{-32}$ | −0.249 |
| standard vs. prioritization | 0.701 | **0.808** | $5.7 \times 10^{-33}$ | −0.266 |
| standard vs. all fixed | 0.701 | **0.826** | $1.9 \times 10^{-34}$ | −0.298 |
| HP only vs. prioritization | 0.799 | **0.808** | 0.265 | −0.024 |
| HP only vs. all fixed | 0.799 | **0.826** | 0.010 | −0.069 |
| prioritization vs. all fixed | 0.808 | **0.826** | 0.143 | −0.044 |
| *Low-priority pharmacophores* | | | | |
| standard vs. HP only | **0.702** | 0.590 | $8.6 \times 10^{-61}$ | +0.344 |
| standard vs. prioritization | 0.702 | **0.739** | $1.6 \times 10^{-6}$ | −0.105 |
| standard vs. all fixed | 0.702 | **0.812** | $7.4 \times 10^{-41}$ | −0.324 |
| HP only vs. prioritization | 0.590 | **0.739** | $2.6 \times 10^{-91}$ | −0.436 |
| HP only vs. all fixed | 0.590 | **0.812** | $5.0 \times 10^{-141}$ | −0.596 |
| prioritization vs. all fixed | 0.739 | **0.812** | $2.0 \times 10^{-19}$ | −0.223 |

##### S2.2.3 Scaffold conditioning

**Table S10**: Pairwise post-relax validity comparisons for scaffold-constrained generation on the MOSES test set. Statistical comparisons are two-sided Mann–Whitney U tests, Holm-corrected within each scaffold definition. Validity is the fraction of samples with a valid post-relaxation molblock per test molecule; higher is better, so a negative Cliff's $\delta$ favors the second-named strategy (B). The better (higher) median in each row is shown in bold.

| Comparison (A vs. B) | Median validity | | $p$ | Cliff's $\delta$ |
|---|---|---|---|---|
| | A | B | | |
| *Random-hetero scaffolds* | | | | |
| Baseline vs. scaffold inpainting | **0.950** | 0.800 | $4.1 \times 10^{-15}$ | +0.645 |
| Baseline vs. scaffold conditioning | **0.950** | 0.950 | 0.158 | +0.111 |
| scaffold conditioning vs. scaffold inpainting | **0.950** | 0.800 | $6.5 \times 10^{-9}$ | +0.480 |
| *BRICS scaffolds* | | | | |
| Baseline vs. scaffold inpainting | **0.950** | 0.950 | 0.966 | −0.003 |
| Baseline vs. scaffold conditioning | 0.950 | **1.000** | 0.022 | −0.206 |
| scaffold conditioning vs. scaffold inpainting | **1.000** | 0.950 | 0.030 | +0.186 |

**Table S11**: Pairwise strain-energy comparisons for scaffold-constrained generation on the MOSES test set. Statistical comparisons are two-sided Mann–Whitney U tests, Holm-corrected within each scaffold definition. Strain energy is reported in kcal/mol; lower is better, so a positive Cliff's $\delta$ favors the second-named strategy (B). The better (lower) median in each row is shown in bold.

| Comparison (A vs. B) | Median strain | | $p$ | Cliff's $\delta$ |
|---|---|---|---|---|
| | A | B | | |
| *Random-hetero scaffolds* | | | | |
| Baseline vs. scaffold inpainting | **0.015** | 0.062 | $5.7 \times 10^{-134}$ | −0.452 |
| Baseline vs. scaffold conditioning | **0.015** | 0.019 | $1.2 \times 10^{-5}$ | −0.080 |
| scaffold conditioning vs. scaffold inpainting | **0.019** | 0.062 | $1.2 \times 10^{-95}$ | −0.381 |
| *BRICS scaffolds* | | | | |
| Baseline vs. scaffold inpainting | **0.015** | 0.017 | 0.055 | −0.035 |
| Baseline vs. scaffold conditioning | 0.015 | **0.009** | $5.0 \times 10^{-54}$ | +0.284 |
| scaffold conditioning vs. scaffold inpainting | **0.009** | 0.017 | $4.2 \times 10^{-53}$ | −0.281 |

###### S2.2.4 Composition

**Table S12**: Mann–Whitney U comparisons of single profile-only conditioning vs. dataset samples, evaluated on the *opposing* similarity metric. Statistical comparisons are two-sided Mann–Whitney U tests, Holm-corrected within each Setting/condition block. Higher value is better; a positive Cliff's $\delta$ favors the first-named strategy (A). The larger median in each row is shown in bold.

| Setting | Comparison (A vs. B) | Median | | $p$ | Cliff's $\delta$ |
|---|---|---|---|---|---|
| | | A | B | | |
| *Similarity to Profile 1* | | | | | |
| Shape | profile 2 only vs. dataset samples | **0.594** | 0.566 | $5.0 \times 10^{-86}$ | +0.184 |
| ESP | profile 2 only vs. dataset samples | **0.565** | 0.556 | $8.0 \times 10^{-10}$ | +0.058 |
| Pharm. | profile 2 only vs. dataset samples | **0.250** | 0.243 | $1.9 \times 10^{-5}$ | +0.040 |
| *Similarity to Profile 2* | | | | | |
| Shape | profile 1 only vs. dataset samples | **0.592** | 0.570 | $1.5 \times 10^{-57}$ | +0.150 |
| ESP | profile 1 only vs. dataset samples | **0.567** | 0.560 | $1.7 \times 10^{-4}$ | +0.035 |
| Pharm. | profile 1 only vs. dataset samples | **0.248** | 0.243 | $3.4 \times 10^{-4}$ | +0.034 |

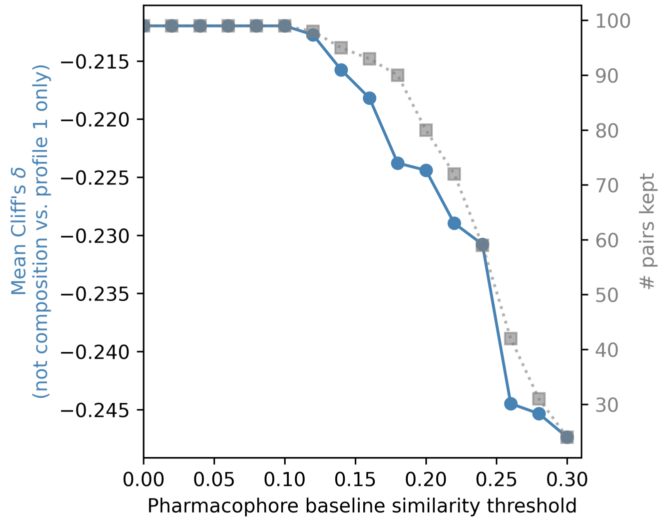

**Fig. S5**: NOT composition threshold sweep. The mean Cliff's $\delta$ when molecule pairs with a pharmacophore similarity less than the threshold are excluded.

**Table S13**: Pairwise Mann–Whitney U comparisons of 3D similarity metrics for AND composition (profile 1 `AND` profile 2) on the MOSES test set, evaluated for similarity to Profile 1 and similarity to Profile 2. Statistical comparisons are two-sided Mann–Whitney U tests, Holm-corrected within each Setting/condition block. Higher similarity is better; a positive Cliff's $\delta$ favors the first-named strategy (A). The larger median in each row is shown in bold.

| Setting | Comparison (A vs. B) | Median | | $p$ | Cliff's $\delta$ |
|---|---|---|---|---|---|
| | | A | B | | |
| *Similarity to Profile 1* | | | | | |
| Shape | and composition vs. profile 1 only | 0.686 | **0.793** | $< 10^{-300}$ | −0.586 |
| ESP | and composition vs. profile 1 only | 0.645 | **0.770** | $< 10^{-300}$ | −0.559 |
| Pharm. | and composition vs. profile 1 only | 0.323 | **0.537** | $< 10^{-300}$ | −0.762 |
| Shape | and composition vs. profile 2 only | **0.686** | 0.594 | $< 10^{-300}$ | +0.598 |
| ESP | and composition vs. profile 2 only | **0.645** | 0.565 | $< 10^{-300}$ | +0.451 |
| Pharm. | and composition vs. profile 2 only | **0.323** | 0.250 | $< 10^{-300}$ | +0.528 |
| Shape | and composition vs. dataset samples | **0.686** | 0.566 | $< 10^{-300}$ | +0.710 |
| ESP | and composition vs. dataset samples | **0.645** | 0.556 | $< 10^{-300}$ | +0.505 |
| Pharm. | and composition vs. dataset samples | **0.323** | 0.243 | $< 10^{-300}$ | +0.549 |
| *Similarity to Profile 2* | | | | | |
| Shape | and composition vs. profile 1 only | **0.683** | 0.592 | $< 10^{-300}$ | +0.606 |
| ESP | and composition vs. profile 1 only | **0.643** | 0.567 | $< 10^{-300}$ | +0.448 |
| Pharm. | and composition vs. profile 1 only | **0.327** | 0.248 | $< 10^{-300}$ | +0.535 |
| Shape | and composition vs. profile 2 only | 0.683 | **0.787** | $< 10^{-300}$ | −0.604 |
| ESP | and composition vs. profile 2 only | 0.643 | **0.763** | $< 10^{-300}$ | −0.567 |
| Pharm. | and composition vs. profile 2 only | 0.327 | **0.529** | $< 10^{-300}$ | −0.759 |
| Shape | and composition vs. dataset samples | **0.683** | 0.570 | $< 10^{-300}$ | +0.699 |
| ESP | and composition vs. dataset samples | **0.643** | 0.560 | $< 10^{-300}$ | +0.487 |
| Pharm. | and composition vs. dataset samples | **0.327** | 0.243 | $< 10^{-300}$ | +0.549 |

**Table S14**: Pairwise Mann–Whitney U comparisons of 3D similarity metrics for AND-NOT composition (profile 1 `AND NOT` profile 2) on the MOSES test set, restricted to the subset of profile pairs whose baseline similarity exceeds 0.3 on all three metrics, evaluated for similarity to Profile 1 and change in similarity to Profile 2 relative to baseline similarity of Profile 1 to Profile 2. Statistical comparisons are two-sided Mann–Whitney U tests, Holm-corrected within each Setting/condition block. Higher similarity (or a greater positive change) is better; a positive Cliff's $\delta$ favors the first-named strategy (A). The larger median in each row is shown in bold.

| Setting | Comparison (A vs. B) | Median | | $p$ | Cliff's $\delta$ |
|---|---|---|---|---|---|
| | | A | B | | |
| *Similarity to Profile 1* | | | | | |
| Shape | not composition vs. profile 1 only | 0.773 | **0.783** | $7.7 \times 10^{-3}$ | $-0.053$ |
| ESP | not composition vs. profile 1 only | 0.719 | **0.761** | $4.4 \times 10^{-13}$ | $-0.145$ |
| Pharm. | not composition vs. profile 1 only | 0.541 | **0.563** | $5.5 \times 10^{-4}$ | $-0.069$ |
| Shape | not composition vs. dataset samples | **0.773** | 0.575 | $< 10^{-300}$ | $+0.852$ |
| ESP | not composition vs. dataset samples | **0.719** | 0.566 | $6.8 \times 10^{-249}$ | $+0.662$ |
| Pharm. | not composition vs. dataset samples | **0.541** | 0.263 | $< 10^{-300}$ | $+0.912$ |
| *Change in Profile Similarity (1→2)* | | | | | |
| Shape | not composition vs. profile 1 only | -0.026 | **-0.004** | $7.5 \times 10^{-33}$ | $-0.240$ |
| ESP | not composition vs. profile 1 only | -0.053 | **-0.014** | $7.2 \times 10^{-55}$ | $-0.313$ |
| Pharm. | not composition vs. profile 1 only | -0.049 | **-0.030** | $7.3 \times 10^{-21}$ | $-0.189$ |
| Shape | not composition vs. dataset samples | **-0.026** | -0.029 | 0.877 | $-0.003$ |
| ESP | not composition vs. dataset samples | -0.053 | **-0.029** | $8.8 \times 10^{-29}$ | $-0.220$ |
| Pharm. | not composition vs. dataset samples | **-0.049** | -0.070 | $3.5 \times 10^{-15}$ | $+0.155$ |

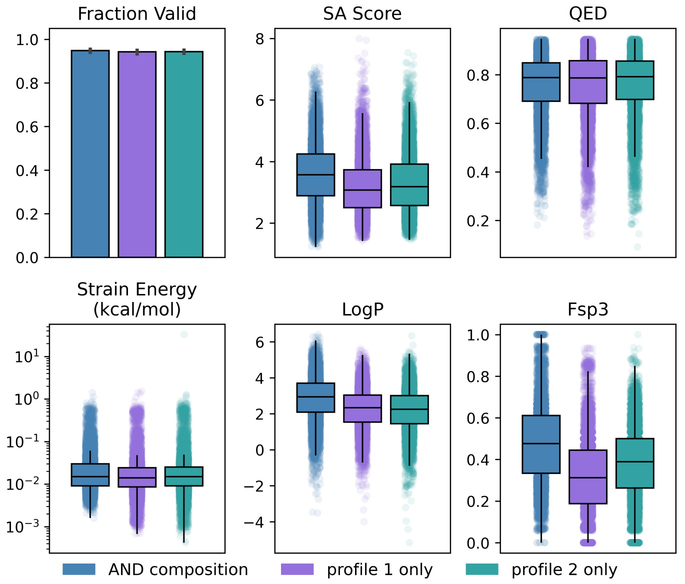

**Fig. S6**: Sample quality of generated samples under AND composition. All metrics aside from strain energy are computed from the post-relaxed state.

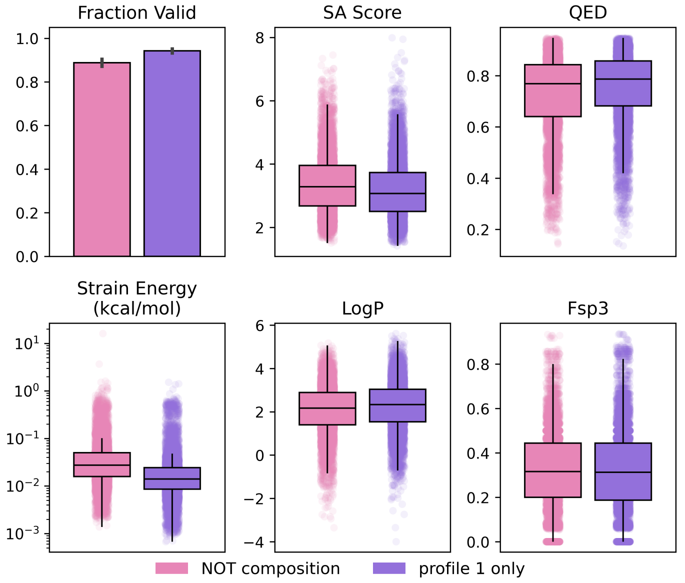

**Fig. S7**: Sample quality of generated samples under NOT composition. All metrics aside from strain energy are computed from the post-relaxed state.

#### S2.3 MolGenBench

We report the full results for the hit-to-lead task of MolGenBench on both ligand- and structure-based settings. Conditioning modalities are summarized as combinations of molecular shape (Sh), electrostatics (E), pharmacophoric features (Ph), pocket atoms (Po), interacting pharmacophores (IPh), interacting atoms (IA), and conserved scaffold (Sc). Metrics assess molecular quality, interaction recovery, docking performance, and diversity (see Sec. 4.5.2 and Suppl. Table S17 for definitions). Models that are statistically indistinguishable share a rank.

**Table S15**: Performance of ligand-based models on the MolGenBench hit-to-lead benchmark. Best and second-best performances are shown in **bold** and underlined, respectively. The mean and 95% confidence intervals are reported.

| | ShEPhERD-2 | FLOWR.ROOT | PhoreGen | OMTRA | CondSemla |
|---|---|---|---|---|---|
| Interaction Info. | Sh+E+Ph | Ph | Sh+Ph | Ph | Ph |
| Conf. Validity ↑ | **0.831**<br>[0.817, 0.845] | 0.712<br>[0.685, 0.737] | 0.290<br>[0.255, 0.327] | 0.727<br>[0.706, 0.745] | 0.785<br>[0.775, 0.795] |
| Diversity ↑ | 0.828<br>[0.824, 0.833] | 0.826<br>[0.816, 0.835] | 0.800<br>[0.796, 0.805] | 0.572<br>[0.557, 0.587] | **0.854**<br>[0.851, 0.856] |
| SA ↑ | 0.647<br>[0.635, 0.658] | 0.666<br>[0.657, 0.676] | 0.719<br>[0.712, 0.725] | 0.730<br>[0.722, 0.738] | **0.825**<br>[0.822, 0.827] |
| QED ↑ | 0.565<br>[0.547, 0.583] | 0.462<br>[0.442, 0.483] | 0.430<br>[0.410, 0.450] | 0.509<br>[0.485, 0.532] | **0.601**<br>[0.592, 0.609] |
| PB Valid ↑ | **0.603**<br>[0.577, 0.630] | 0.062<br>[0.048, 0.077] | 0.046<br>[0.037, 0.056] | 0.572<br>[0.547, 0.595] | 0.323<br>[0.292, 0.354] |
| PB Energy Ratio ↑ | **0.871**<br>[0.858, 0.883] | 0.814<br>[0.794, 0.833] | 0.504<br>[0.454, 0.556] | 0.776<br>[0.757, 0.793] | 0.836<br>[0.828, 0.844] |
| Int. Score >p50 ↑ | 0.321<br>[0.280, 0.363] | 0.145<br>[0.116, 0.177] | 0.192<br>[0.151, 0.235] | **0.398**<br>[0.351, 0.445] | 0.099<br>[0.081, 0.119] |
| Vina<p50 ↑ | **0.317**<br>[0.265, 0.371] | 0.162<br>[0.128, 0.196] | 0.201<br>[0.163, 0.241] | 0.260<br>[0.213, 0.309] | 0.123<br>[0.088, 0.163] |
| Mean Vina ↓ | **-8.59**<br>[-8.81, -8.36] | -7.57<br>[-7.76, -7.38] | -7.95<br>[-8.14, -7.75] | -8.30<br>[-8.50, -8.10] | -7.54<br>[-7.69, -7.40] |
| Min Vina ↓ | **-12.06**<br>[-12.35, -11.75] | -11.10<br>[-11.36, -10.83] | -11.26<br>[-11.54, -10.98] | -10.89<br>[-11.16, -10.62] | -10.23<br>[-10.42, -10.04] |
| Ref. Tanimoto ↓ | 0.199<br>[0.191, 0.207] | 0.174<br>[0.163, 0.186] | 0.132<br>[0.126, 0.138] | 0.451<br>[0.432, 0.470] | **0.110**<br>[0.108, 0.112] |
| Int. Diversity ↑ | 0.756<br>[0.745, 0.767] | 0.739<br>[0.719, 0.758] | 0.662<br>[0.644, 0.678] | 0.417<br>[0.396, 0.438] | **0.808**<br>[0.800, 0.816] |

**Table S16**: Performance of pocket-based models on the MolGenBench hit-to-lead benchmark. Best and second-best performances are shown in **bold** and underlined, respectively. The mean and 95% confidence intervals are reported. Ranks are computed including the ShEPhERD–2 model (Sh+E+Ph+IA) and are slightly different from those in the main text.

| | ShEPhERD–2 | ShEPhERD–2 | FLOWR.ROOT | PhoreGen | OMTRA | DiffDec | DiffSBDD |
|---|---|---|---|---|---|---|---|
| Interaction Info. | Sh+E+IPh | Sh+E+Ph+IA | IA+Po | Sh+IPh | Ph+Po | Sc+Po | Sc+Po |
| Conf. Validity ↑ | **0.836** [0.822, 0.849] | 0.721 [0.693, 0.747] | **0.835** [0.816, 0.853] | 0.352 [0.305, 0.400] | 0.729 [0.709, 0.748] | 0.485 [0.450, 0.521] | 0.659 [0.630, 0.686] |
| Diversity ↑ | **0.813** [0.806, 0.819] | 0.761 [0.749, 0.773] | 0.782 [0.772, 0.791] | **0.809** [0.799, 0.818] | 0.662 [0.653, 0.672] | 0.567 [0.550, 0.583] | 0.696 [0.670, 0.721] |
| SA ↑ | 0.654 [0.642, 0.665] | 0.656 [0.643, 0.668] | 0.695 [0.682, 0.707] | 0.696 [0.685, 0.707] | **0.734** [0.725, 0.742] | **0.741** [0.727, 0.753] | 0.701 [0.689, 0.712] |
| QED ↑ | 0.553 [0.533, 0.572] | 0.532 [0.511, 0.553] | 0.465 [0.443, 0.488] | 0.481 [0.454, 0.507] | 0.453 [0.428, 0.477] | **0.605** [0.584, 0.626] | 0.559 [0.543, 0.575] |
| PB Valid ↑ | 0.639 [0.614, 0.663] | 0.594 [0.567, 0.621] | **0.777** [0.754, 0.798] | 0.290 [0.248, 0.334] | 0.661 [0.640, 0.680] | 0.466 [0.433, 0.501] | 0.559 [0.526, 0.592] |
| PB Energy Ratio ↑ | **0.879** [0.867, 0.889] | 0.776 [0.749, 0.802] | **0.885** [0.872, 0.898] | 0.532 [0.468, 0.597] | 0.819 [0.802, 0.836] | 0.547 [0.512, 0.582] | 0.700 [0.671, 0.728] |
| Int. Score >p50 ↑ | 0.393 [0.348, 0.440] | **0.514** [0.456, 0.572] | **0.525** [0.476, 0.574] | 0.368 [0.311, 0.426] | 0.457 [0.406, 0.509] | 0.374 [0.319, 0.431] | 0.331 [0.288, 0.376] |
| Vina<p50 ↑ | 0.320 [0.267, 0.374] | 0.328 [0.275, 0.384] | **0.425** [0.359, 0.491] | 0.263 [0.211, 0.318] | 0.268 [0.221, 0.317] | 0.190 [0.151, 0.233] | 0.171 [0.135, 0.209] |
| Mean Vina ↓ | -8.61 [-8.83, -8.38] | -8.66 [-8.89, -8.43] | **-9.11** [-9.35, -8.87] | -8.33 [-8.58, -8.09] | -8.35 [-8.57, -8.12] | -7.91 [-8.10, -7.72] | -7.76 [-7.96, -7.56] |
| Min Vina ↓ | -12.08 [-12.39, -11.76] | -11.93 [-12.26, -11.61] | **-12.83** [-13.19, -12.48] | -11.70 [-12.11, -11.29] | -11.55 [-11.85, -11.25] | -10.66 [-10.93, -10.40] | -11.22 [-11.49, -10.94] |
| Ref. Tanimoto ↓ | 0.222 [0.212, 0.232] | 0.288 [0.273, 0.303] | 0.184 [0.173, 0.195] | **0.130** [0.121, 0.140] | 0.420 [0.406, 0.434] | 0.329 [0.314, 0.345] | 0.231 [0.217, 0.246] |
| Int. Diversity ↑ | **0.714** [0.700, 0.727] | 0.606 [0.582, 0.628] | 0.647 [0.625, 0.668] | 0.676 [0.657, 0.695] | 0.506 [0.487, 0.524] | 0.427 [0.398, 0.456] | 0.620 [0.596, 0.644] |

**Table S17**: Definitions of evaluation metrics used in our analysis of the MolGenBench benchmark. PoseBusters metrics were computed using the PoseBusters benchmark suite, docking scores using AutoDock Vina, interaction metrics using ProLIF, and molecular similarity metrics using RDKit Morgan fingerprints.

| Category | Metric | Definition | Better |
|---|---|---|---|
| Molecular quality | Conformer Validity | Fraction of generated molecules passing PoseBusters conformer validity checks, which assess the chemical plausibility of the generated three-dimensional structure independent of the protein environment. | ↑ |
| | Diversity | Mean pairwise Tanimoto distance between generated molecules using Morgan fingerprints. | ↑ |
| | SA | Synthetic accessibility (SA) score [33]. | ↑ |
| | QED | Quantitative estimate of drug-likeness [34]. | ↑ |
| Protein compatibility | PB Valid | Fraction of generated protein–ligand complexes passing all PoseBusters validation checks, including ligand geometry and protein–ligand interaction constraints [27]. | ↑ |
| | PB Energy Ratio | Mean PoseBusters energy-ratio score, which estimates conformational strain by comparing the generated conformer with 50 conformers embedded with ETKDG and relaxed with MMFF94. | ↑ |
| Interaction recovery | Interaction Score >p50 | Fraction of generated molecules exhibiting at least the median number of conserved protein–ligand interactions observed among known active compounds for the corresponding target series. Interactions are identified using ProLIF. | ↑ |
| Docking | Vina <p50 | Fraction of generated molecules with an AutoDock Vina score better than the median score of known active compounds for the corresponding target. | ↑ |
| | Mean Vina | Mean AutoDock Vina docking score across generated molecules. | ↓ |
| | Min Vina | Best (lowest) AutoDock Vina docking score obtained among generated molecules for each target, averaged across targets. | ↓ |
| Novelty | Reference Tanimoto | Morgan fingerprint Tanimoto similarity between each generated molecule and its conditioning reference ligand. | ↓ |
| | Interaction Diversity | Diversity of interaction-mediating molecular substructures identified by ProLIF, computed using fingerprint-based similarity. | ↑ |

#### S2.4 Case Studies

##### S2.4.1 Bioisosteric fragment merging

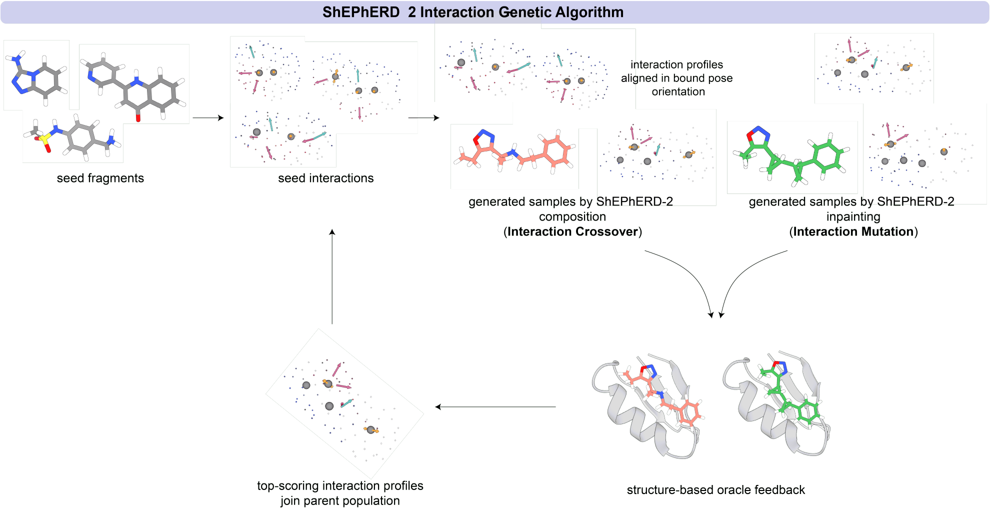

**Fig. S8**: Interaction-aware genetic algorithm derives seed interaction profiles from bound poses of input molecules. New interaction profiles are generated by composition or inpainting, and the associated generated molecules are evaluated by structure-based oracles, which output a score and a pose that inform subsequent selection.

**Table S18**: Top-$N$ docking score and molecular-property statistics (mean ± std across 3 replicate runs) for interaction-space optimization vs. a 2D pharmacophore fingerprint-based bioisosteric merging baseline [35]. The baseline library was bootstrapped to match the number of molecules generated per optimization run, so that Top-N is comparable.

| | Top-1 | | Top-10 | | Top-50 | |
|---|---|---|---|---|---|---|
| Metric | Exploitation | Baseline | Exploitation | Baseline | Exploitation | Baseline |
| Docking Score | -8.66 ± 0.23 | -7.58 ± 0.04 | -8.16 ± 0.02 | -7.45 ± 0.03 | -7.78 ± 0.03 | -7.15 ± 0.04 |
| QED | 0.62 ± 0.14 | 0.51 ± 0.05 | 0.64 ± 0.07 | 0.54 ± 0.04 | 0.63 ± 0.07 | 0.61 ± 0.02 |
| cLogP | 4.35 ± 2.40 | 4.64 ± 0.49 | 4.78 ± 0.79 | 3.42 ± 0.08 | 4.79 ± 0.64 | 2.99 ± 0.03 |
| MW (Da) | 431.3 ± 18.1 | 401.1 ± 26.6 | 421.2 ± 4.2 | 410.2 ± 12.6 | 416.2 ± 4.6 | 394.8 ± 3.8 |
| SA Score | 4.00 ± 0.24 | 2.13 ± 0.07 | 4.07 ± 0.10 | 2.50 ± 0.07 | 4.07 ± 0.03 | 2.50 ± 0.04 |
| TPSA ($Å^2$) | 58.3 ± 30.6 | 80.0 ± 9.4 | 57.2 ± 15.3 | 93.6 ± 4.3 | 57.3 ± 14.6 | 91.4 ± 2.4 |
| Rotatable Bonds | 3.7 ± 1.2 | 5.0 ± 0.0 | 4.7 ± 0.3 | 5.3 ± 0.4 | 5.2 ± 0.2 | 4.9 ± 0.2 |
| $F_{sp^3}$ | 0.54 ± 0.17 | 0.08 ± 0.05 | 0.56 ± 0.12 | 0.19 ± 0.03 | 0.56 ± 0.14 | 0.19 ± 0.01 |

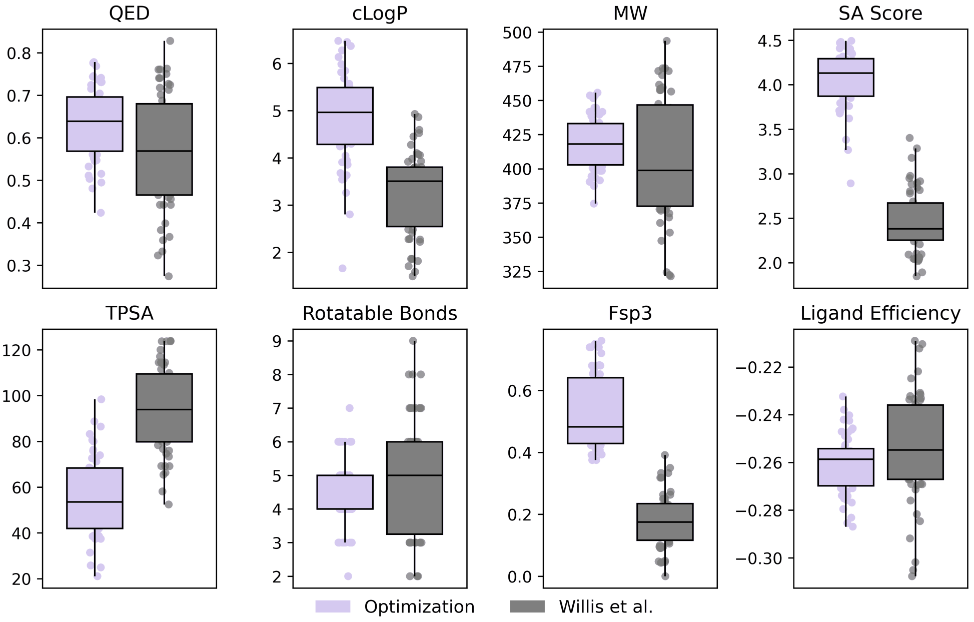

**Fig. S9**: Comparison of molecular properties of the pooled top-50 molecules between ShEPhERD–2 and a 2D pharmacophore fingerprint-based bioisosteric merging baseline [35].

**Table S19**: Top-$N$ Boltz-2-predicted affinity ($\log_{10}$ $IC_{50}$/µM) and molecular-property statistics (mean ± std across 3 replicate runs) for interaction-space optimization vs. a 2D pharmacophore fingerprint-based bioisosteric merging baseline [35], both scored with the Boltz-2 affinity oracle. The baseline library was bootstrapped to match the number of molecules generated per optimization run, so that Top-N is comparable.

| | Top-1 | | Top-10 | | Top-50 | |
|---|---|---|---|---|---|---|
| Metric | Exploitation | Baseline | Exploitation | Baseline | Exploitation | Baseline |
| Affinity (log µM) | -0.67 ± 0.03 | 0.24 ± 0.10 | -0.48 ± 0.11 | 0.50 ± 0.04 | -0.21 ± 0.12 | 0.75 ± 0.03 |
| QED | 0.72 ± 0.08 | 0.50 ± 0.03 | 0.74 ± 0.12 | 0.59 ± 0.05 | 0.74 ± 0.09 | 0.67 ± 0.01 |
| cLogP | 4.91 ± 0.49 | 4.85 ± 0.26 | 4.64 ± 0.16 | 3.57 ± 0.04 | 4.50 ± 0.34 | 3.12 ± 0.11 |
| MW (Da) | 339.6 ± 90.2 | 422.9 ± 4.7 | 338.0 ± 66.4 | 388.8 ± 18.4 | 341.5 ± 65.4 | 368.1 ± 2.1 |
| SA Score | 4.01 ± 0.42 | 2.14 ± 0.14 | 3.99 ± 0.16 | 2.60 ± 0.06 | 3.99 ± 0.27 | 2.68 ± 0.06 |
| TPSA (Å$^2$) | 35.3 ± 8.4 | 71.2 ± 1.8 | 44.4 ± 14.8 | 76.1 ± 3.2 | 46.0 ± 13.1 | 73.5 ± 2.2 |
| Rotatable Bonds | 5.0 ± 1.7 | 6.7 ± 0.6 | 4.5 ± 1.3 | 6.2 ± 0.5 | 4.5 ± 0.6 | 5.9 ± 0.1 |
| $F_{sp^3}$ | 0.56 ± 0.06 | 0.26 ± 0.04 | 0.56 ± 0.01 | 0.34 ± 0.02 | 0.57 ± 0.02 | 0.39 ± 0.02 |

**Table S20**: Top-$N$ docking score and molecular-property statistics (mean ± std across 3 replicate runs) for pharmacophore-conditioned interaction-space optimization vs. zero-shot fragment aggregate profile generation [11].

| | Top-1 | | Top-10 | | Top-50 | |
|---|---|---|---|---|---|---|
| Metric | Exploitation | Baseline | Exploitation | Baseline | Exploitation | Baseline |
| Docking Score | -8.83 ± 0.17 | -8.61 ± 0.05 | -8.49 ± 0.20 | -8.38 ± 0.03 | -8.17 ± 0.22 | -8.06 ± 0.02 |
| QED | 0.44 ± 0.09 | 0.35 ± 0.09 | 0.42 ± 0.03 | 0.35 ± 0.02 | 0.44 ± 0.02 | 0.37 ± 0.02 |
| cLogP | 4.66 ± 1.91 | 5.27 ± 1.03 | 3.82 ± 1.16 | 4.56 ± 0.03 | 3.62 ± 0.52 | 4.18 ± 0.22 |
| MW (Da) | 446.8 ± 24.1 | 468.9 ± 7.8 | 432.9 ± 20.5 | 478.1 ± 10.4 | 427.5 ± 9.1 | 467.8 ± 1.2 |
| SA Score | 4.03 ± 0.16 | 4.12 ± 0.08 | 3.85 ± 0.39 | 4.14 ± 0.10 | 3.78 ± 0.28 | 4.08 ± 0.06 |
| TPSA (Å$^2$) | 100.9 ± 24.8 | 84.9 ± 22.5 | 98.4 ± 15.1 | 99.1 ± 2.2 | 101.5 ± 12.9 | 100.2 ± 1.9 |
| Rotatable Bonds | 3.3 ± 0.6 | 6.0 ± 1.0 | 4.0 ± 1.2 | 5.7 ± 0.2 | 4.4 ± 0.8 | 5.7 ± 0.1 |
| $F_{sp^3}$ | 0.27 ± 0.13 | 0.29 ± 0.11 | 0.21 ± 0.11 | 0.28 ± 0.01 | 0.21 ± 0.09 | 0.27 ± 0.01 |

##### S2.4.2 Dual target and selectivity design

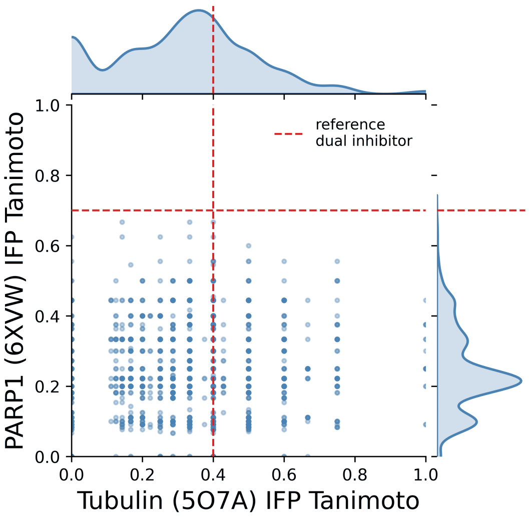

**Fig. S10**: Joint ProLIF interaction fingerprint similarity (IFP Tanimoto) to reference Tubulin and PARP1 inhibitors across AND composition samples. Reference dual inhibitor joint interaction fingerprint similarities are indicated by the red dashed lines.

##### S2.4.3 Modality hopping

**Table S21**: One-sample $z$-test of the reference peptidomimetic's drug-like molecular properties against the corresponding ShEPhERD–2 generated-sample distribution. Results tabulated per macrocyclic peptide mimicry target with a reported uncorrected two-sided $p$-value from a standard Gaussian distribution. $^{*}p < 0.05$, $^{**}p < 0.01$, $^{***}p < 0.001$, and not significant $p$-values are bolded.

| Target | Property | Sample mean | Sample SD | Reference | $z$ | $p$-value | |
|---|---|---|---|---|---|---|---|
| HCMV | QED | 0.303 | 0.113 | 0.249 | −0.48 | **0.634** | |
| HCMV | cLogP | 3.125 | 1.656 | 5.032 | 1.15 | **0.249** | |
| HCMV | MW | 493.977 | 32.728 | 629.721 | 4.15 | $3.4 \times 10^{-5}$ | *** |
| HCMV | SA Score | 4.627 | 0.699 | 4.018 | −0.87 | **0.384** | |
| HCMV | TPSA | 94.537 | 26.527 | 125.330 | 1.16 | **0.246** | |
| HCMV | Rotatable Bonds | 12.257 | 2.339 | 8.000 | −1.82 | **0.069** | |
| HCMV | $F_{sp^3}$ | 0.506 | 0.081 | 0.250 | −3.15 | 0.002 | ** |
| NNMT | QED | 0.376 | 0.169 | 0.259 | −0.69 | **0.490** | |
| NNMT | cLogP | 2.580 | 1.789 | 6.011 | 1.92 | **0.055** | |
| NNMT | MW | 411.551 | 54.109 | 564.093 | 2.82 | 0.005 | ** |
| NNMT | SA Score | 4.450 | 0.837 | 3.883 | −0.68 | **0.498** | |
| NNMT | TPSA | 80.259 | 25.373 | 97.760 | 0.69 | **0.490** | |
| NNMT | Rotatable Bonds | 8.898 | 2.001 | 6.000 | −1.45 | **0.148** | |
| NNMT | $F_{sp^3}$ | 0.603 | 0.155 | 0.219 | −2.47 | 0.013 | * |

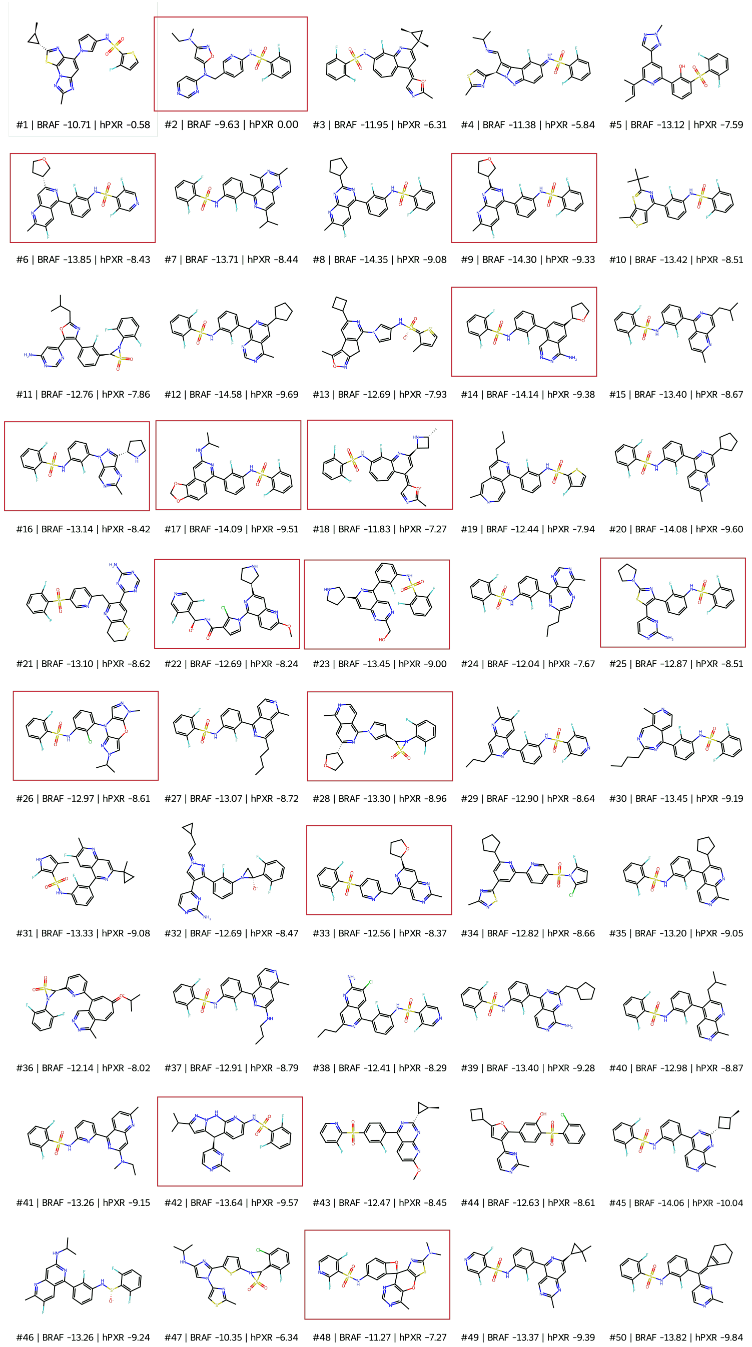

**Fig. S11**: Top 50 generated samples by highest predicted selectivity for BRAF over hPXR. Highlighted samples (red) mirror the same selectivity strategy experimentally confirmed in [36] with dabrafenib's tert-butyl replaced with a large polar substituent.

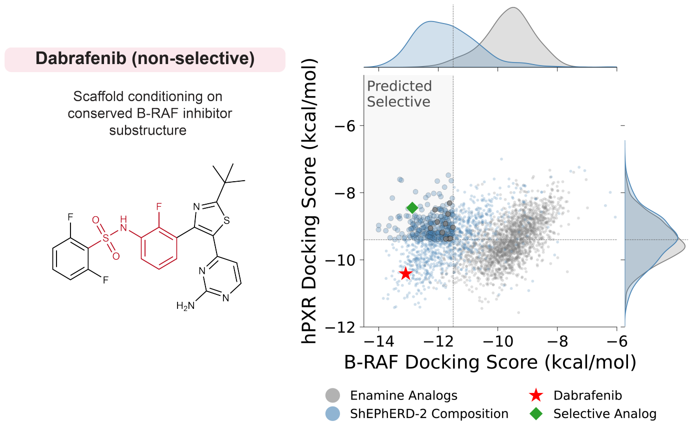

**Fig. S12**: Distributions of BRAF and hPXR docking scores for NOT composition dabrafenib redesigns using scaffold conditioning. The conditioned scaffold (red) is a conserved substructure across BRAF inhibitors.

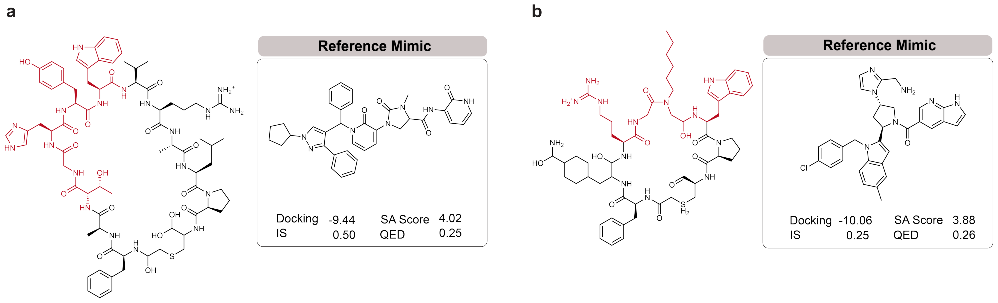

**Fig. S13**: Macrocyclic peptide binders of **(a)** human cytomegalovirus protease and **(b)** nicotinamide N-methyltransferase, with interface areas highlighted in red, alongside their associated reference small-molecule mimics.

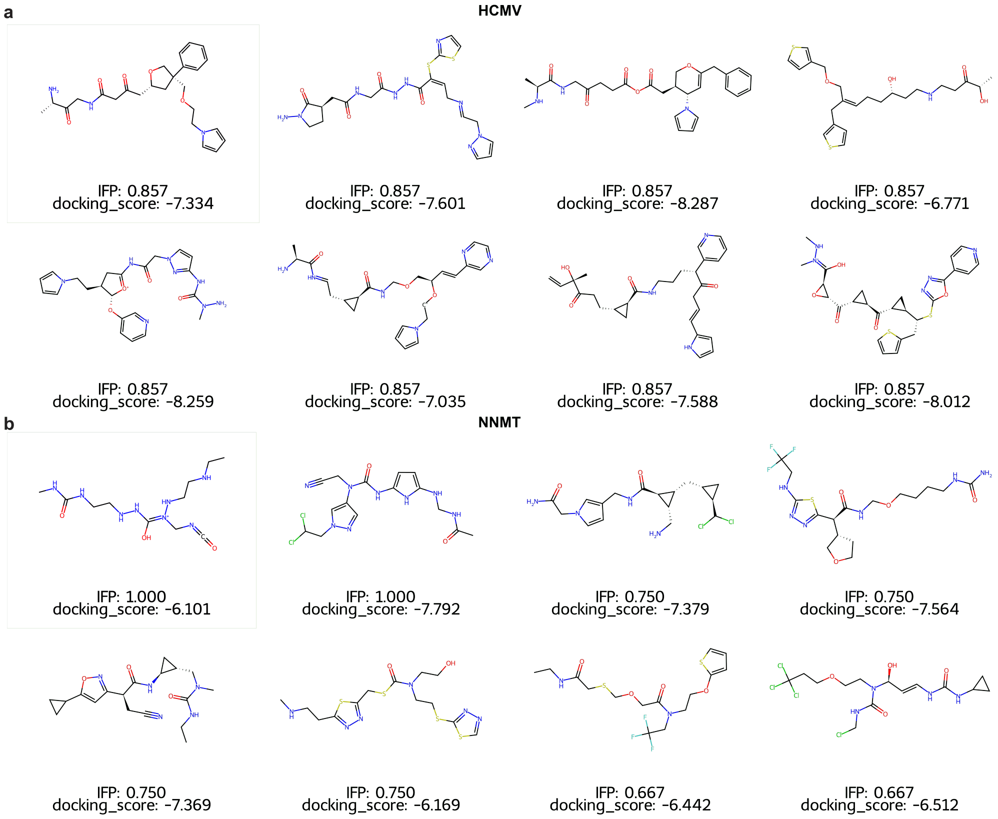

**Fig. S14**: Top generated molecules by interaction similarity (IFP) to interface peptide interactions for **(a)** human cytomegalovirus protease and **(b)** nicotinamide N-methyltransferase.

### S3 Derivations

#### S3.1 Composition Formulation

##### S3.1.1 AND Composition

**Notation.** We simplify the notation used in Suppl. Sec. S1.2-S1.4 for readability. Let $x := \boldsymbol{X}_G^{(\sigma)}$ be the generated structure at the current noise level $\sigma = \sigma_i$, and let $x' := \boldsymbol{X}_G^{(\sigma')}$ be the next state at $\sigma' = \sigma_{i-1}$. Write $p(x) := p_{\text{data}} * \mathcal{N}(0, \sigma^2 I)$ for the marginal density of the noised profile, and $p(\cdot \mid c)$ for the corresponding conditional marginal given a target profile $c = \boldsymbol{X}_K^c$. Let $D(\cdot\,; \sigma)$ be the denoiser at level $\sigma$; by Tweedie's identity, it realizes the score of $p$, $\nabla_x \log p(x) = \big(D(x;\sigma) - x\big)/\sigma^2$.

**Definition 1** (Sampler) The score form of the predict–renoise update (S10) is

$$x' \;=\; x + \sigma^2\, \nabla_x \log p(x) + \sigma'\, \boldsymbol{\epsilon}, \qquad \boldsymbol{\epsilon} \sim \mathcal{N}(\mathbf{0}, \boldsymbol{I}), \tag{S22}$$

where $\nabla_x \log p(x)$ may be the score of any (tilted) marginal we supply.

**Lemma 1** (Denoiser form of the sampler) For a single (un)conditional marginal, (S22) reduces to

$$x' \;=\; D(x;\sigma) + \sigma'\, \boldsymbol{\epsilon}, \tag{S23}$$

and $x' = D^{c_i}(x;\sigma) + \sigma' \boldsymbol{\epsilon}$ in the conditional case.

*Proof* Insert $\nabla_x \log p(x) = (D(x;\sigma) - x)/\sigma^2$ into (S22); the $\sigma^2$ factors cancel and the $x$ terms telescope:

$$x' = x + (D(x;\sigma) - x) + \sigma' \boldsymbol{\epsilon} = D(x;\sigma) + \sigma' \boldsymbol{\epsilon}. \qquad \square$$

**Assumption 1** (Conditional independence of profiles) Given the noised sample $x$, the profiles are conditionally independent: $p(c_1, \ldots, c_n \mid x) = \prod_{i=1}^n p(c_i \mid x)$.

**Proposition 1** (AND composition) *Under Assumption 1, the sampler (S22) driven by the (guidance-weighted) joint score of $p(x \mid c_1, \ldots, c_n)$ is*

$$\boxed{x' = D(x;\sigma) + \sum_{i=1}^{n} w_{c_i} \big(D^{c_i}(x;\sigma) - D(x;\sigma)\big) + \sigma_{i-1}\, \boldsymbol{\epsilon}}$$

*where $D^{c_i}(x;\sigma) = D(x;\sigma,c_i)$ is the denoiser conditioned on a profile via inpainting.*

Intuition: *start from the unconditional prediction and add each profile's pull away from the unconditional prediction.*

*Proof* Under Assumption 1, Bayes' rule gives the target as a product of experts

$$p(x \mid c_1, \ldots, c_n) \;\propto\; p(x,\, c_1, \ldots, c_n) \;=\; p(x) \prod_{i=1}^{n} p(c_i \mid x), \tag{S24}$$

and, again by Bayes' rule and dropping the $x$-independent prior $p(c_i)$,

$$p(c_i \mid x) \;\propto\; \frac{p(x \mid c_i)}{p(x)}. \tag{S25}$$

Taking logarithms of (S24) and inserting (S25), all $x$-independent terms collect into constant $C$:

$$\log p(x \mid c_1, \ldots, c_n) = \log p(x) + \sum_{i=1}^{n} \Big( \log p(x \mid c_i) - \log p(x) \Big) + C. \tag{S26}$$

Differentiating (S26) with respect to $x$ kills the constant $C$:

$$\nabla_x \log p(x \mid c_1, \ldots, c_n) = \nabla_x \log p(x) + \sum_{i=1}^{n} \Big( \nabla_x \log p(x \mid c_i) - \nabla_x \log p(x) \Big). \tag{S27}$$

Now we apply the Tweedie identity to each score term, using the unconditional denoiser $D$ for $p(x)$ and the target profile-conditional denoiser $D^{c_i}$ for $p(x \mid c_i)$:

$$\nabla_x \log p(x \mid c_1, \ldots, c_n) = \frac{D(x;\sigma) - x}{\sigma^2} + \sum_{i=1}^{n} \left( \frac{D^{c_i}(x;\sigma) - x}{\sigma^2} - \frac{D(x;\sigma) - x}{\sigma^2} \right). \tag{S28}$$

Next we introduce guidance weights to control the influence of each target interaction profile. We refer to the gradient of the tilted density of the joint conditional $p(x \mid c_1, \ldots, c_n)$ with guidance weights $w_i$ as $\nabla_x \log \tilde{p}$.

$$\nabla_x \log \tilde{p} = \frac{D - x}{\sigma^2} + \sum_{i=1}^{n} w_i \left( \frac{D^{c_i} - x}{\sigma^2} - \frac{D - x}{\sigma^2} \right) = \frac{D - x}{\sigma^2} + \sum_{i=1}^{n} w_i \, \frac{D^{c_i} - D}{\sigma^2}, \tag{S29}$$

where setting $w_i = 1$ recovers the exact product of experts in (S28).

Insert (S29) into the reverse step (S22):

$$\begin{aligned} x' &= x + \sigma^2 \left[ \frac{D - x}{\sigma^2} + \sum_{i=1}^{n} w_i \, \frac{D^{c_i} - D}{\sigma^2} \right] + \sigma' \epsilon \\ &= x + (D - x) + \sum_{i=1}^{n} w_i (D^{c_i} - D) + \sigma' \epsilon \\ &= D(x;\sigma) + \sum_{i=1}^{n} w_i \big( D^{c_i}(x;\sigma) - D(x;\sigma) \big) + \sigma' \epsilon. \end{aligned} \tag{S30}$$

□

###### S3.1.2 NOT Composition

We now express one desired *on-target* interaction profile while *suppressing* a set of undesired *off-target* profiles $\mathrm{off}_1, \ldots, \mathrm{off}_n$. Mirroring the AND construction, we tilt the on-target marginal by the ratio of on- to off-target likelihoods, with guidance weights $w_i$; we again write $\tilde{p}$ for the resulting tilted density:

$$\tilde{p}(x) \;\propto\; p(x \mid \mathrm{on}) \prod_{i=1}^{n} \left( \frac{p(x \mid \mathrm{on})}{p(x \mid \mathrm{off}_i)} \right)^{w_i}. \tag{S31}$$

**Proposition 2** (NOT composition) *The sampler* (S22) *driven by the score* $\nabla_x \log \tilde{p}$ *of the tilted target* (S31) *is*

$$\boxed{x' = D^{\text{on}}(x;\sigma) + \sum_{i=1}^{n} w_i \Big( D^{\text{on}}(x;\sigma) - D^{\text{off}_i}(x;\sigma) \Big) + \sigma' \epsilon}$$

Intuition*: start from the on-target prediction and, for each off-target profile, add a pull* away *from that profile.*

*Proof* This is the AND template of Proposition 1 under the substitutions $D \mapsto D^{\text{on}}$ (the base becomes the on-target prediction) and, in each summand, $D^{c_i} \mapsto D^{\text{on}}$, $D \mapsto D^{\text{off}_i}$. We give the steps explicitly.

Taking logarithms of (S31) and differentiating, every $x$-independent normalizer vanishes:

$$\nabla_x \log \tilde{p}(x) = \nabla_x \log p(x \mid \text{on}) + \sum_{i=1}^{n} w_i \Big( \nabla_x \log p(x \mid \text{on}) - \nabla_x \log p(x \mid \text{off}_i) \Big). \tag{S32}$$

Applying Tweedie's identity to each score term, with the on-target denoiser $D^{\text{on}}$ for $p(x \mid \text{on})$ and the off-target denoisers $D^{\text{off}_i}$ for $p(x \mid \text{off}_i)$, the $-x$ terms cancel inside each difference:

$$\nabla_x \log \tilde{p}(x) = \frac{D^{\text{on}} - x}{\sigma^2} + \sum_{i=1}^{n} w_i \frac{D^{\text{on}} - D^{\text{off}_i}}{\sigma^2}. \tag{S33}$$

Inserting (S33) into the reverse step (S22), exactly as in the AND case, the $\sigma^2$ factors cancel and the $x$ terms telescope:

$$\begin{aligned} x' &= x + \sigma^2 \left[ \frac{D^{\text{on}} - x}{\sigma^2} + \sum_{i=1}^{n} w_i \frac{D^{\text{on}} - D^{\text{off}_i}}{\sigma^2} \right] + \sigma' \epsilon \\ &= D^{\text{on}}(x;\sigma) + \sum_{i=1}^{n} w_i \big( D^{\text{on}}(x;\sigma) - D^{\text{off}_i}(x;\sigma) \big) + \sigma' \epsilon. \end{aligned} \tag{S34}$$

□
